# *Klebsiella pneumoniae* type VI secretion system-mediated microbial competition is PhoPQ controlled and reactive oxygen species dependent

**DOI:** 10.1101/698415

**Authors:** Daniel Storey, Alan McNally, Mia Åstrand, Joana Sá-Pessoa Graca Santos, Isabel Rodriguez-Escudero, Bronagh Elmore, Leyre Palacios, Helina Marshall, Laura Hobley, Maria Molina Martin, Victor J. Cid, Tiina A. Salminen, Jose A. Bengoechea

**Author notes:** School of Biosciences, University of Nottingham, Leicestershire, UK.

## Abstract

*Klebsiella pneumoniae* is recognized as an urgent threat to human health due to the increasing isolation of multidrug resistant strains. Hypervirulent strains are a major concern due to their ability to cause life-threating infections in healthy hosts. The type VI secretion system (T6SS) is widely implicated in microbial antagonism, and it mediates interactions with host eukaryotic cells in some cases. In silico search for genes orthologous to T6SS component genes and T6SS effector genes across 700 *K. pneumoniae* genomes shows extensive diversity in T6SS genes across the *K. pneumoniae* species. Temperature, oxygen tension, pH, osmolarity, iron levels, and NaCl regulate the expression of the T6SS encoded by a hypervirulent *K. pneumoniae* strain. Polymyxins and human defensin 3 also increase the activity of the T6SS. A screen for regulators governing T6SS uncover the correlation between the transcription of the T6SS and the ability to kill *E. coli* prey. Whereas H-NS represses the T6SS, PhoPQ, PmrAB, Hfq, Fur, RpoS and RpoN positively regulate the T6SS. *K. pneumoniae* T6SS mediates intra and inter species bacterial competition. This antagonism is only evident when the prey possess an active T6SS. The PhoPQ two component system governs the activation of *K. pneumoniae* T6SS in bacterial competitions. Mechanistically, PhoQ periplasmic domain, and the acid patch within, is essential to activate *K. pneumoniae* T6SS. *Klebsiella* T6SS also mediates anti-fungal competition. We have delineated the contribution of each of the individual VgrGs in microbial competition, and identified VgrG4 as a T6SS effector. Structurally, domain DUF2345 of VgrG4 is sufficient to intoxicate bacteria and yeast. ROS generation mediates the antibacterial effects of VgrG4, and the antitoxin Sel1E protects against the toxic activity of VgrG4. Our findings provide a better understanding of the regulation of the T6SS in bacterial competitions, and place ROS as an early event in microbial competition.

**AUTHOR SUMMARY:** *Klebsiella pneumoniae* has been singled out as an “urgent threat to human health” due to extremely drug resistant strains. Numerous studies investigate the molecular mechanisms underlying antibiotic resistance in *K. pneumoniae*, while others dissect the virulence strategies of this pathogen. However, there is still limited knowledge on the fitness of *Klebsiella* in the environment, and particularly the competition of *Klebsiella* with other species. Here, we demonstrated that *Klebsiella* exploits the type VI secretion system (T6SS) nanoweapon to kill bacterial competitors and fungi. *K. pneumoniae* perceives T6SS attacks from bacterial competitors, resulting in retaliation against the aggressive cell. The perception of the attack involved the sensor PhoPQ and led to the up-regulation of the T6SS. We identified one of the toxins deployed by the T6SS to antagonize other microbes, and revealed how *Klebsiella* protects itself from this toxin. Our findings provide a better understanding of the T6SS role in microbial competition and uncover new aspects on how bacteria regulate T6SS-mediated microbial antagonism.

## INTRODUCTION

The type VI secretion system (T6SS) is a bacterial nanomachinery that delivers substrates into a target cell in an one-step process.The system was first described as a cluster of impaired in nitrogen fixation (*imp*) genes in *Rhizobium leguminosarum* [1]. Shortly after, Rao and co-workers identified the T6SS of *Edwardsiella tarda* in a mass spectrometric screen for secreted virulence factors [2]. Initial studies reported that T6SSs contributed to virulence and/or interaction with eukaryotic cells in multiple bacterial species, primarily pathogens (*Vibrio cholerae, Burkholderia ssp., Pseudomonas aeruginosa, E. tarda, Salmonella gallinarum*) [3-6]. However, substantial experimental data argue for its broad significance in bacterial fitness in the environment. The first antibacterial T6SS was described in *P. aeruginosa* in 2010 [7], and since then the T6SS has been implicated in efficient killing of competitors, quorum sensing and horizontal gene transfer [8-11].

T6SS are encoded within clusters which all contain 13 conserved core genes named *tssA-M* encoding the proteins making up the basic secretion apparatus [12-15]. ClpV/TssH is a cytoplasmic AAA+ ATPase allowing recycling of the T6SS machinery. Additional copies of essential core components, *tssD*/*hcp* and *tssI*/*vgrG*, can be found outside the main T6SS cluster and are often associated with genes encoding putative effector proteins. Bioinformatics and structural evidence sustain the notion that the T6SS apparatus is closely related to components of a contractile bacteriophage forming a cell puncture device [12-15]. The T6SS inner tube is composed of hexamers of Hcp, and it is believed that VgrG sits at the top of the Hcp tube. PAAR domain containing proteins have been shown to bind the distal end of VgrG to complete the tip of the machinery but also to act as a platform for recruitment of effectors [16]. The evidence indicates that an effector can either be fused to a homologue of one of the components of the Hcp-VgrG-PAAR structure as an additional domain, or can interact non-covalently with one of the components of this structure [17]. The first characterized effector domain was the C-terminal actin cross-linking domain of the VgrG-1 protein secreted by the T6SS of *V. cholerae* [18]. Many other effectors are encoded in proximity to *vgrG, hcp* or *paar* genes strongly suggesting that their secretion is associated with the neighbouring core component; this has been extensively investigated in the case of *Serratia marcescens* and *P. aeruginosa*.

*Klebsiella pneumoniae* has been singled out as an “urgent threat to human health” by the World Health Organization due to extremely drug resistant strains. More than a third of the *K. pneumoniae* isolates reported to the European Centre for Disease Prevention and Control were resistant to at least one antimicrobial group, being the most common resistance phenotype the combined resistance to fluoroquinolones, third-generation cephalosporins and aminoglycosides. Numerous studies investigate the molecular mechanisms underlying antibiotic resistance in *K. pneumoniae*, while others dissect the virulence strategies of this pathogen [19,20]. However, there is still limited knowledge on the fitness of *Klebsiella* in the environment, and particularly the competition of *Klebsiella* with other species. In this context, a first analysis of four *Klebsiella* genomes revealed that this pathogen may encode a T6SS with the 13 core *T6SS* genes organized in two syntenic segments [21].Three putative VgrGs and two Hcp were described as potential effector proteins through their sequence similarities to *V. cholerae* and *P. aeruginosa* effector proteins [21]. Our recent comparative genomics study indicates that the T6SS is part of the *Klebsiella* core genome [22], and two different phospholipases, PLD1 and Tle1, have been identified as T6SS effectors [22,23].

In the present study, we characterize the T6SS encoded by a hypervirulent *K. pneunomiae* strain and investigate its role in intra and inter bacterial species, and anti-fungal competition. Our findings demonstrate that interaction with competing bacteria expressing T6SS triggers the activation of the PhoPQ two-component system leading to increased transcription of the T6SS and killing of competitor bacteria. Moreover, polymyxins, last line antibiotic against multidrug *Klebsiella* infections, and human defensins also induce T6SS-dependent killing in a PhoPQ dependent manner. In this work, we also identify the antibacterial effector domain fused to VgrG4 and demonstrate that its toxicity is through the generation of reactive oxygen species (ROS), highlighting the role of ROS in antibacterial competition.

## RESULTS

### Organization of the T6SS gene clusters in *K. pneumoniae* CIP52.145

Strain *K. pneumoniae* CIP52.145 (hereafter Kp52145) belongs to the *K. pneumoniae* KpI group which includes the vast majority of strains associated with human infection, including numerous multidrug-resistant and hypervirulent clones [24]. Kp52145 encodes all virulence functions significantly associated with invasive community-acquired disease in humans [22,24]. Kp52145 possess three T6SS loci (Fig 1A). Locus I encodes all the known conserved core genes encoding the proteins making up the basic secretion apparatus including ClpV/TssH, Hcp1/TssD1, and VgrG1/TssI1. A putative effector is found downstream of *vgrG1*. In contrast, locus II is incomplete, *hcp*, *clpV*, *vipA/tssB*, and *vipB/tssC* are missing and there is an insertion in *sciN/tssJ* (Fig 1A). However, it does contain another *vgrG*, we termed *vgrG2*, and three putative effectors, locus tag BN49_0937, BN49_0942, BN49_0943, the latter one containing a characteristic PAAR domain. Two putative immunity proteins, locus tag BN49_0941 and BN49_0936, are also found in this cluster. As we have recently described [22], locus III is also incomplete and contains an insertion of nine proteins including another *vgrG*, we termed *vgrG4*, a PLD1 effector, and a PAAR protein (Fig 1A). We have recently demonstrated that *pld1* encodes a virulence factor essential in the pneumonia mouse model [22]. There are no reported studies of the function of the proteins encoded by the five genes containing *sel1* repeats in *K. pneumoniae*, although in *Legionella pneumophila* similar Sel1 containing proteins have been implicated in interactions with host cells [25]. In silico analysis of the Kp52145 genome revealed the presence of three additional putative *hcp* genes, locus tag BN49_1035, BN49_2747 and BN49_2887, and a cluster of five genes, locus tag BN49_4908 to BN49_4913, encoding four putative effectors and a PAAR protein (Fig 1B).

**FIGURE 1.**
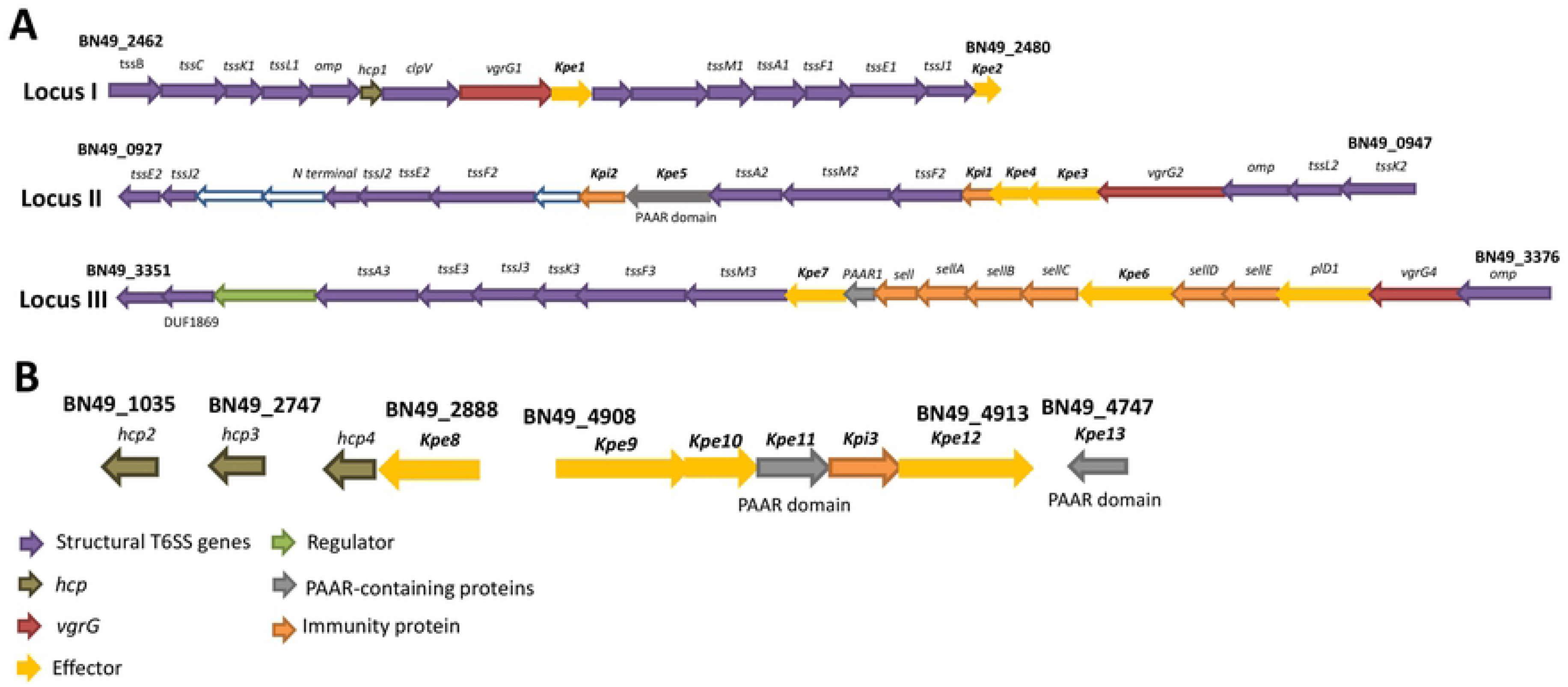
Predicted T6SS gene loci *K. pneumoniae* CIP52.145. (A) Analysis of the three gene loci encoding the T6SS. Locus I encodes all the known conserved core genes, whereas loci II and III are incomplete. Each loci is characterized by a distinct *vgrG* and a different set of putative effector(s) and immunity proteins. (B) Additional T6SS related genes found elsewhere in the genome. In both panels the colour code of the genes indicate putative structural genes, *hcp*, *vgrG*, effector (denoted as Kpe for *<u>K</u>. <u>p</u>neumoniae* <u>e</u>ffectors), immunity protein (denoted as Kpi for *<u>K</u>. <u>p</u>neumoniae* immunity protein), regulator, and PAAR-domain containing protein.

### Diversity of T6SS locus in *K. pneumoniae*

We sought to investigate the diversity of the T6SS locus in different *K. pneumoniae* strains. Initially, we focused on strains ATCC43816, widely used in virulence studies, NTUH-K2044 and SGH10, isolates of the clonal group CG23 causing liver abscesses [26,27], NJST258_1, isolate of the clonal group ST258 [28], and strain MGH78578, the first *K. pneumoniae* multidrug resistant isolate sequenced. To analyse these genomes we used the integrated database SecReT6 (http://db-mml.sjtu.edu.cn/SecReT6/). Locus I is conserved in all strains, and it contains all the conserved genes known to be essential to build a T6SS (S1 Fig). *vgrG1* is found in all strains in locus I. Notably, there is a significant diversity in the number of effectors and immunity proteins present in this locus, and every strain analysed has a different repertoire. In all strains, locus II is also incomplete, and this locus is also characterized by the presence of a *vgrG*. Likewise Kp52.145, MGH75878, encodes *vgrG2* in this locus whereas ATCC431816 and NTUH-K2044 encode *vgrG4*, and SGH10 and NJST258_1 encode a new *vgrG* we termed *vgrG3*. Each of the strains analysed encode also different repertoire of putative effectors in this locus. Only ATCC43816 and MGH75878 present a third T6SS locus, being different to that encoded by Kp52145. Collectively these findings demonstrate that locus I encodes a complete T6SS although there is considerable genome diversity in terms of effectors, *vgrGs*, and number of T6SS loci.

To obtain a global picture of the T6SS in *K. pneumoniae*, we downloaded 700 *K. pneumoniae* genomes from NCBI, selected to represent the broad phylogenetic structure of the species. Using the SecRet6 database we searched for genes orthologous to T6SS component genes (Fig 2A) and T6SS effector genes (Fig 2B) across the genome data set. Our analysis shows extensive diversity in T6SS genes across the *K. pneumoniae* species, with most of this diversity exhibited as high levels of sequence dissimilarity rather than loss of genes. This diversity is independent of population structure which is indicative of frequent and extensive recombination in the *Klebsiella* T6SS locus, analogous to that observed in capsule encoding regions [29]. To determine the potential diversity present in the key *vgrG1* gene, we created a pangenome of the downloaded genomes and blasted all ∼65,000 genes for orthologous sequences of the *vgrG1* gene of Kp52145. Our analysis identified 33 different orthologous *vgrG1* genes within the pangenome, suggesting that extensive T6SS diversity occurs at all levels from component genes through to the essential-for-function *vgrG1*.

**FIGURE 2.**
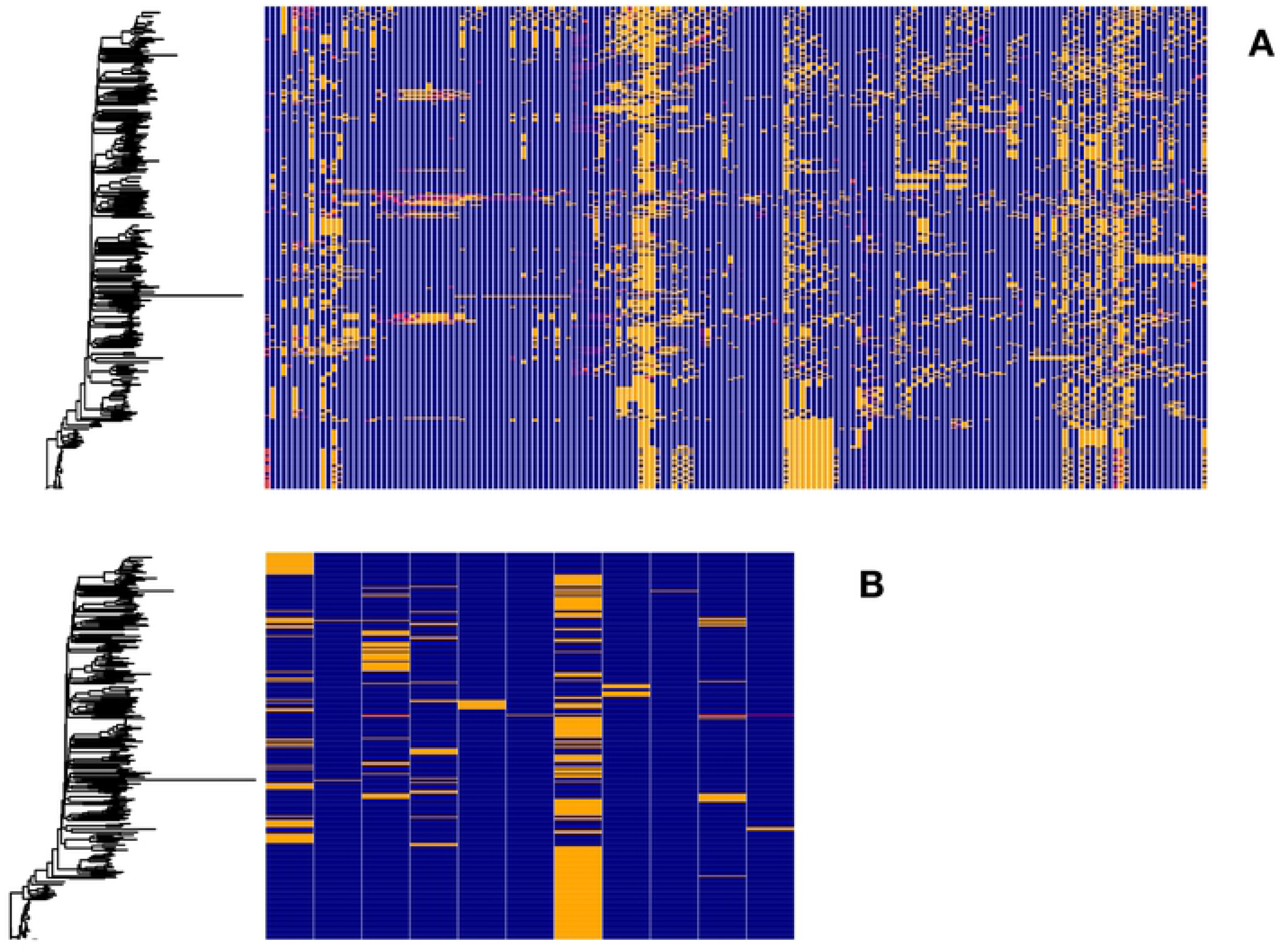
Diversity of *K. pneumoniae* T6SS. Diversity of T6SS component (A) and effector (B) genes across 500 *K. Pneumoniae* genomes. Each horizontal block represents an individual gene present in the SecRet6 database. Presence of the gene at greater than 80% nucleotide diversity is represented by a blue block. Absence, indicated by lack of an orthologous gene sequence at greater than 25% nucleotide identity is presented by a red block. Orthologous sequences at a nucleotide identity between 25% and 80% are represented by yellow blocks.

### Kp52145 does not outcompete *Escherichia coli* in a T6SS-dependent manner in Lysogeny broth

The T6SS of different pathogens has been shown to mediate killing of other bacteria. To determine whether Kp52145 exhibits T6SS-mediated antibacterial activity, we used strain *E. coli* MG1655 harbouring a plasmid conferring resistance to apramycin as prey in bacterial killing assays (Table 1). Co-incubation of *E coli* with Kp52145 showed no difference in killing of *E. coli*, while a dramatic reduction was observed when *E. coli* was incubated with *S. marcescens* Db10 (S2 Fig). In an attempt to understand the lack of antibacterial effect of Kp52145 T6SS, we hypothesized that the *Klebsiella* capsule polysaccharide (CPS) interferes with the contact-dependent killing. We then tested the *cps*-deficient strain 52145-Δ*manC* for killing of *E. coli*. However, the *cps* mutant did not kill *E. coli* in these conditions (S2 Fig), suggesting that the presence of CPS does not interfere with T6SS-dependent killing.

**Table 1.**
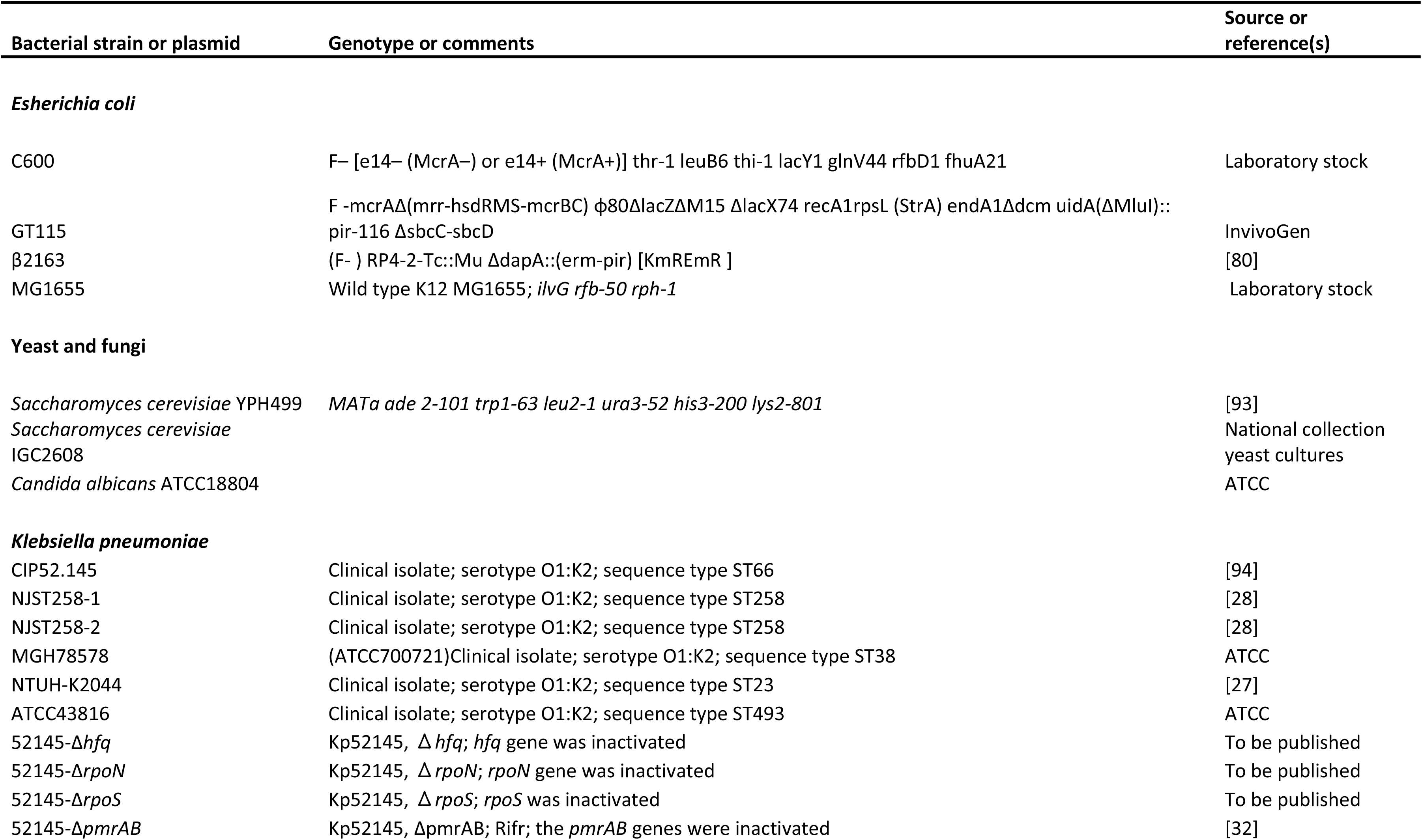
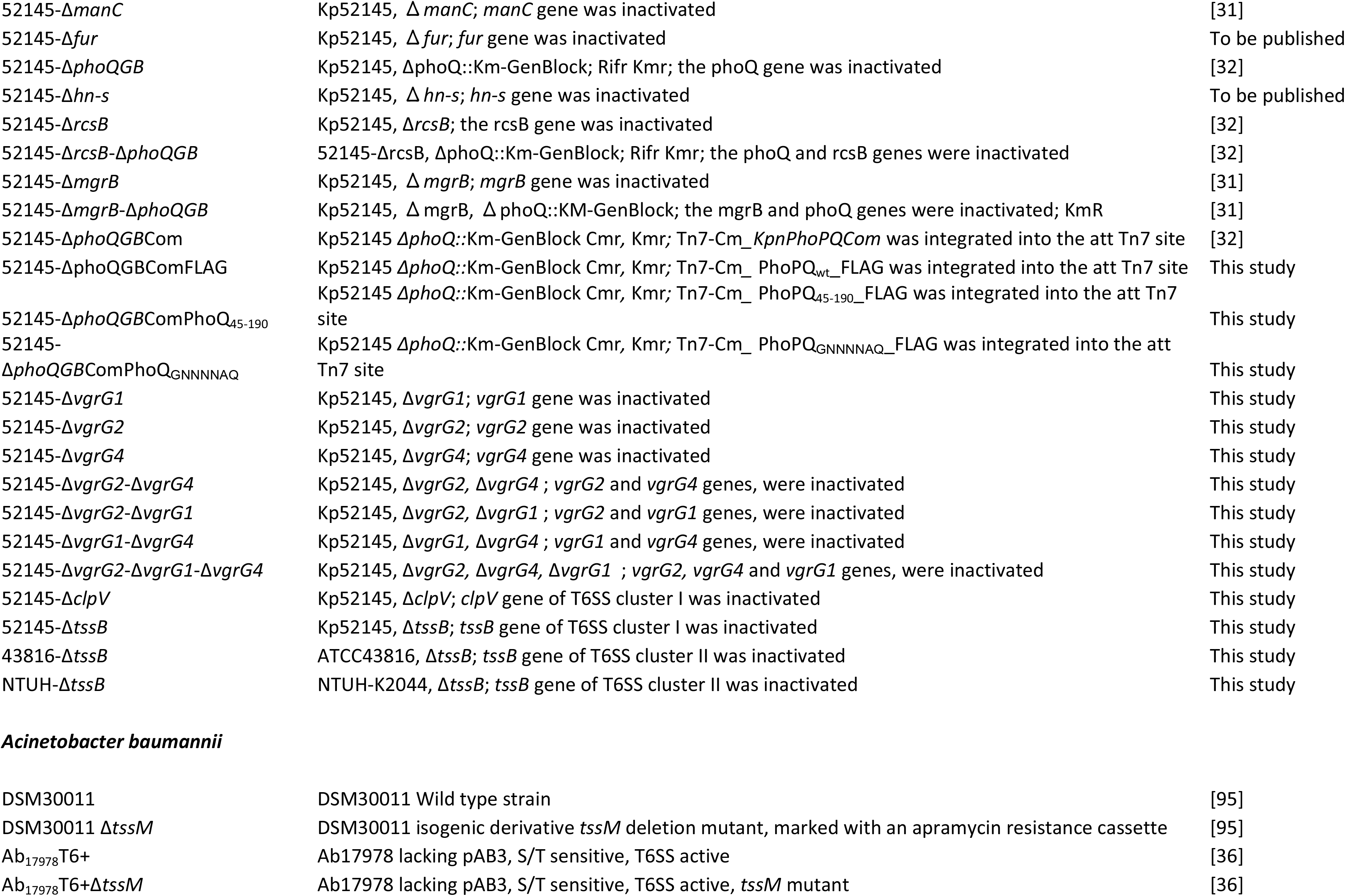
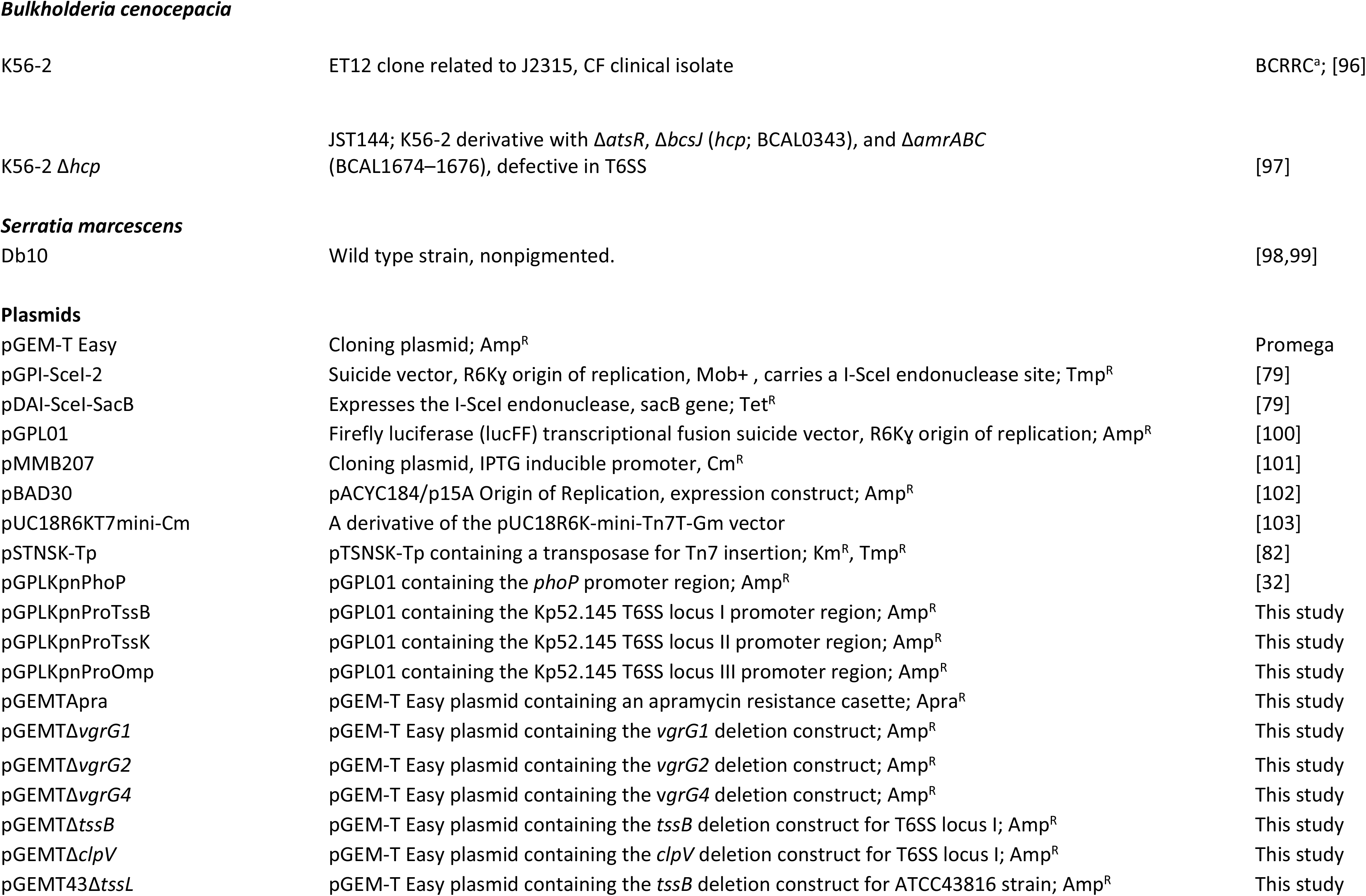
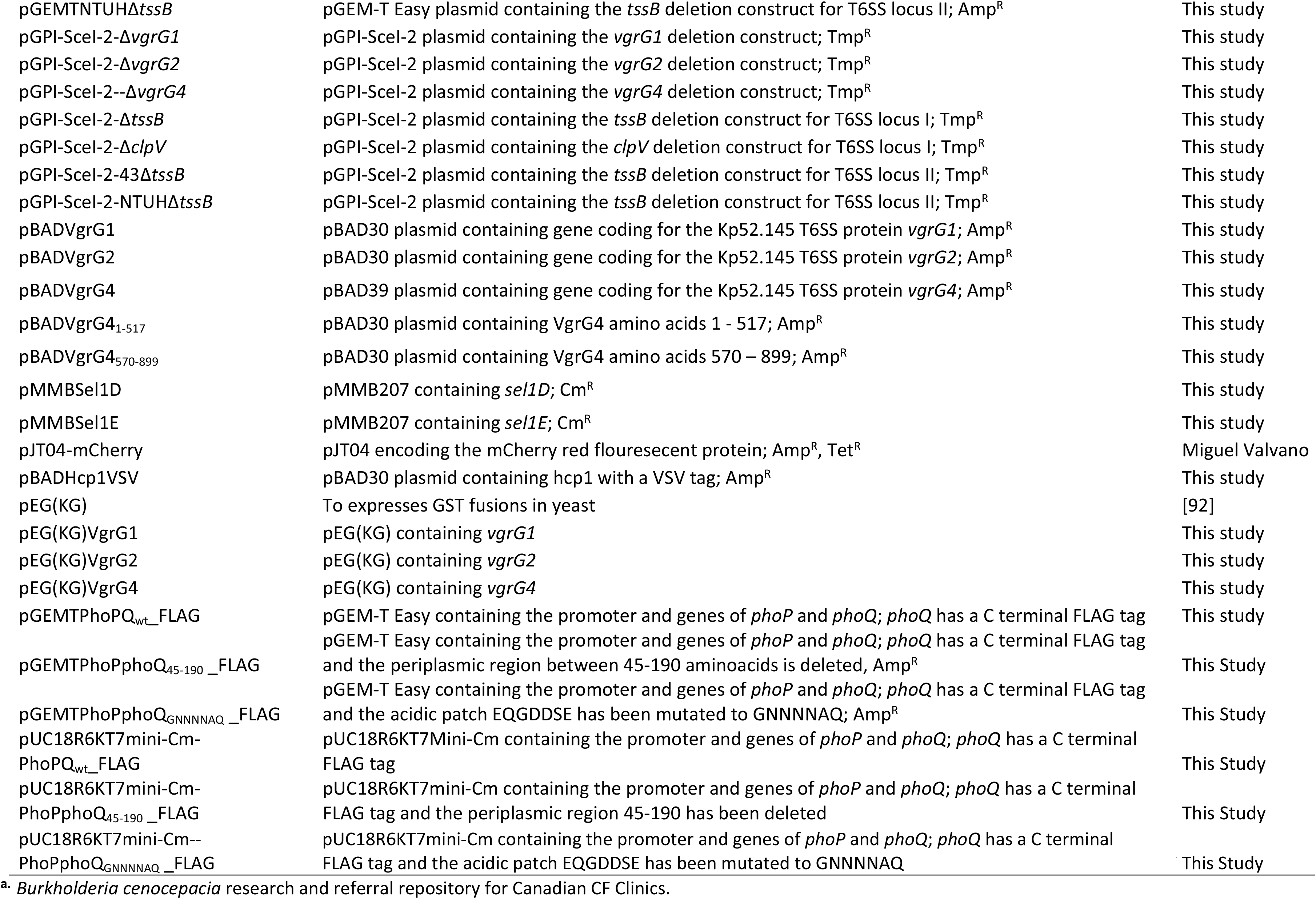
Strains and plasmids used in this study.

### Environmental signals affecting *K. pneumoniae* T6SS expression

To explain the lack of T6SS-dependent killing of Kp52145 when bacteria are grown in lysogeny broth (LB) we speculated that the expression of Kp52145 T6SS is regulated by environmental signals in the hope of identifying conditions in which the system could be up-regulated. To investigate the T6SS transcriptional pattern, we constructed one reporter strain carrying nondisruptive chromosomal transcriptional fusion containing a promoterless luciferase firelfly gene (*lucFF*) under the control of the promoter region of the first gene of the T6SS gene locus I, *tssB::lucFF.* We focused on locus I because this is the conserved one in *Klebsiella* strains, and it encodes all the essential components known to be necessary to assemble a T6SS. We first investigated whether growth phase may affect the pattern of expression of the *tssB::lucFF* fusion. Supplementary Fig 3A shows that the highest levels of luciferase activity were observed when the reporter strain reached mid-exponential phase. Therefore, luciferase activity was measured at mid-exponential phase in subsequent experiments testing different environmental conditions. We next tested the impact of temperature, pH, oxygen tension, osmolarity and ionic strength, and iron levels on the expression of the transcriptional fusion. Results are shown as heat-map indicating when the luciferase activity peaked (Fig 3A). Experiments examining the impact of temperature established that luciferase activity was higher at 37^0^C than at 20°C (Fig 3A), and, therefore, the subsequent experiments testing other conditions were performed growing the reporter strain at 37°C. *K. pneumoniae* survives over a wide range of pH (pH 4.0 to 10.0). Luciferase activity was assayed in cultures grown in different buffered media inoculated in parallel. In the range of pH tested, luciferase activity peaked at pH 6.0 (Fig 3A). To get an indication whether T6SS expression might be regulated in response to oxygen tension, we assessed the effect of aeration using 5-ml cultures grown under the following conditions of increasing aeration: in a 15-ml tube incubated vertically shaking at 180 rpm (condition 1), in a 15-ml slanted tube (45°) shaking at 180 rpm (condition 2), in a 25-ml tube incubated vertically shaking at 180 rpm (condition 3), and in a 30-ml tube incubated vertically shaking at 180 rpm (condition 4) (Fig 3A). Vigorous aeration significantly reduced the luciferase levels compared to the 5-ml standing tube (condition 1), in which the oxygen tension is probably low (Fig 3A). Iron restriction conditions were achieved by growing the reporter strain in medium with increasing concentrations of the iron chelator 2-2’ bipyridyl. Luciferase levels peaked when bacteria were grown with the highest concentration of the iron chelator (Fig 3A), suggesting that low iron concentrations up-regulate the expression of the locus I of the T6SS. The effects of ionic strength versus osmolarity were tested in LB with increasing concentrations of salt (0 to 595 mM KCl) or rhamnose and sucrose. The levels of luciferase activity decreased similarly with increasing concentrations of the solutes, supporting that high osmolarity down-regulates the expression of this T6SS locus (Fig 3A). However, high levels of NaCl did not have an inhibitory effect, and, in fact, we observed a significant increase in the luciferase activity when bacteria were grown in LB containing 595 mM NaCl (Fig 3A), suggesting that the environment signal triggered by NaCl is the sodium concentration.

**FIGURE 3.**
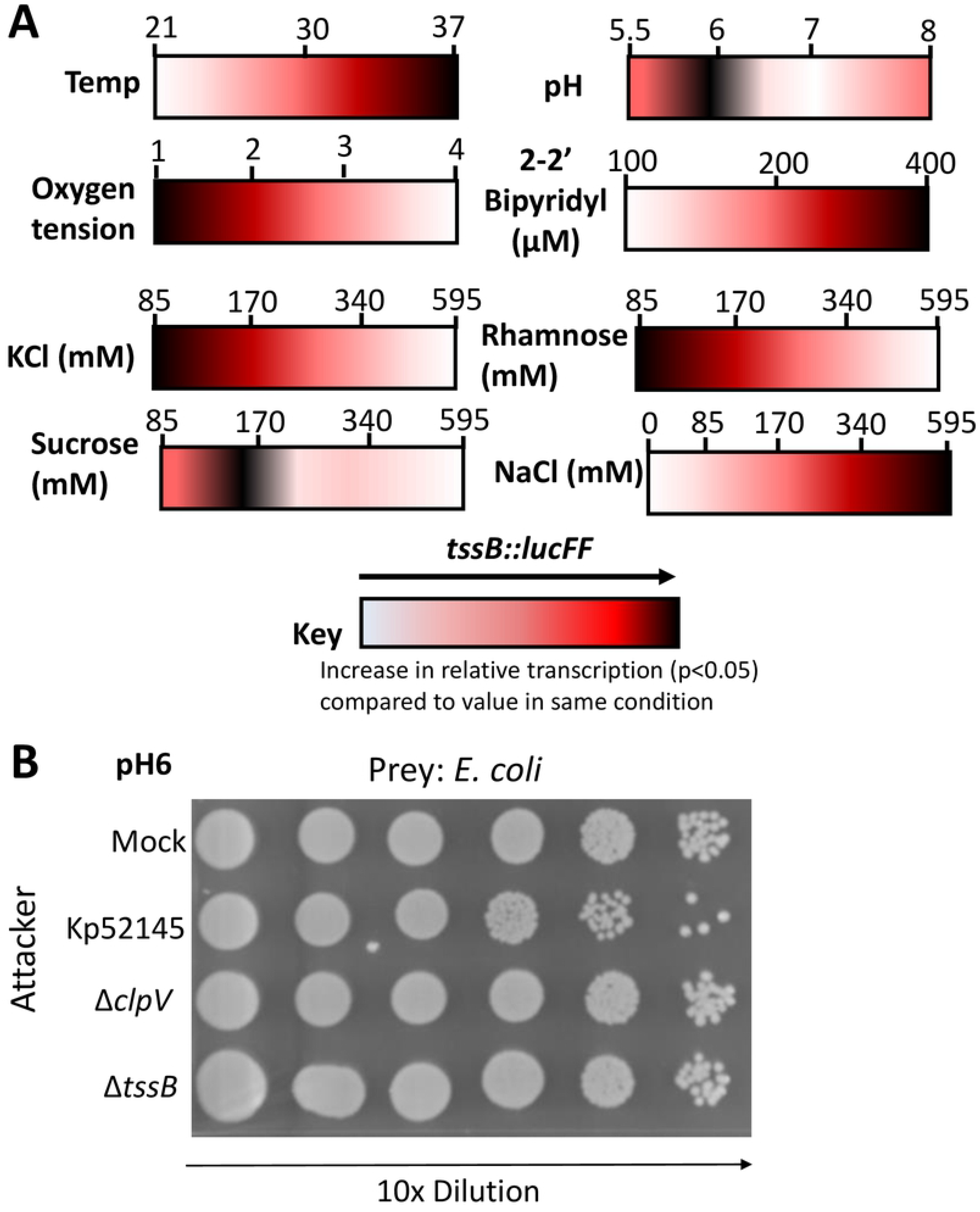
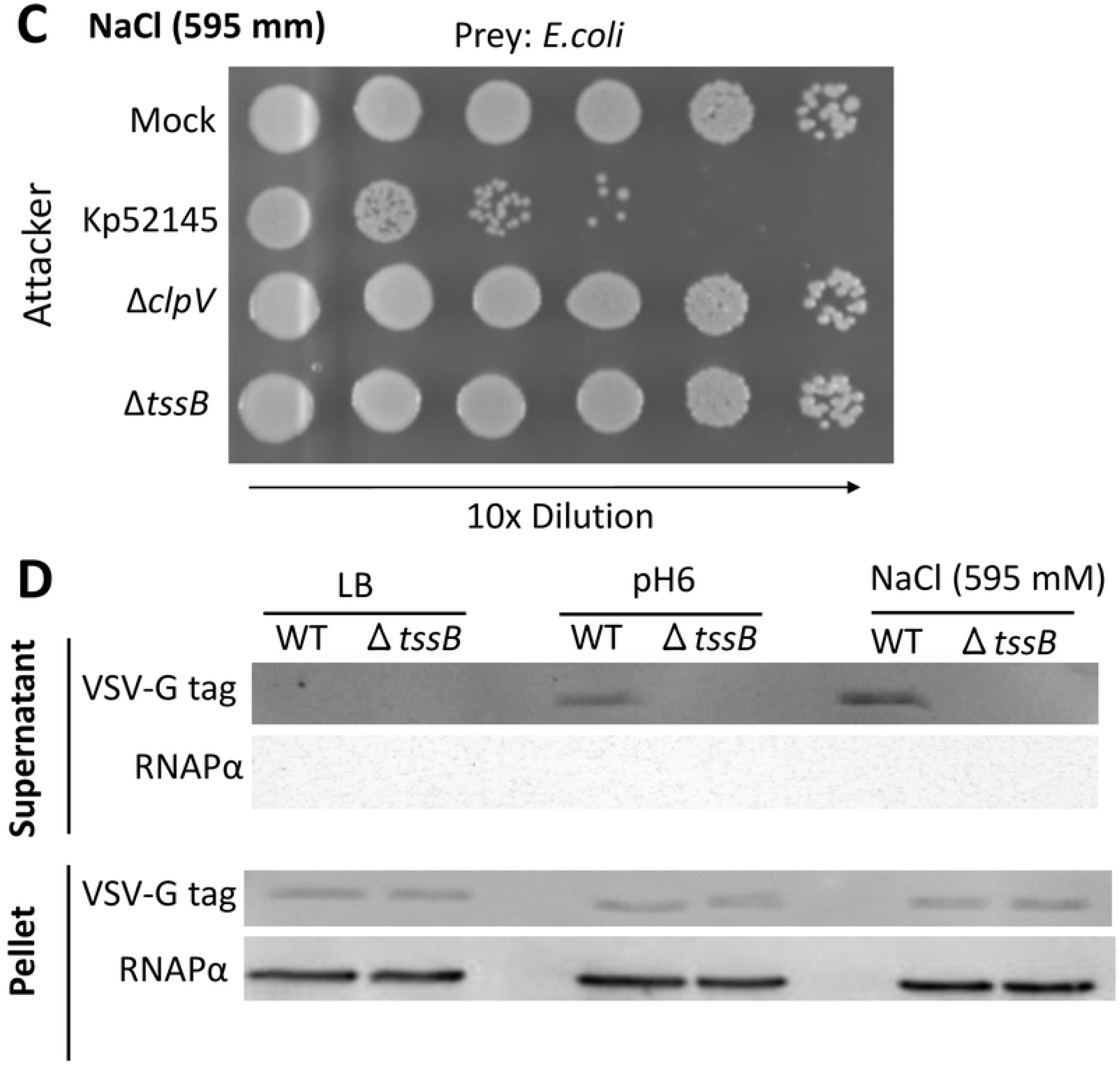
Regulation of *K. pneumoniae* T6SS. (A) Heat-map analysis of the luciferase levels of the transcriptional fusion *tssB::lucFF* in the background of the wild-type strain grown until exponential phase at different temperatures (Temp); pH (LB was buffered as follows: 100 mM MES pH 5.5 and 6, HEPES pH7, MOPS pH8), oxygen tension (1, 15-ml tube incubated vertically shaking at 180 rpm; 2, 15-ml slanted tube (45°) shaking at 180 rpm; 3, 25-ml tube incubated vertically shaking at 180 rpm; 4, 30-ml tube incubated vertically shaking at 180 rpm), and different concentrations of KCl, rhamnose, sucrose, NaCl and the iron chelator 2-2’ bipyridyl. (B, C) Bacterial killing mediated by *K. pneumoniae* CIP52.145 (Kp52145), 52145-Δ*clpV* (Δ*clpV*), 52145-Δ*tssB* (Δ*tssB*) against *E. coli* MG1655. After 6 h incubation in LB_pH6_, LB_NaCl_ and bacteria were recovered, diluted in PBS 0 to 10^-5^ and spotted in LB agar plate. (D) Western blot analysis using an anti-VSV-G antibody demonstrating the presence of Hcp1 in the supernatants and cell pellets of *K. pneumoniae* CIP52.145 (Kp52145), and 52145-Δ*tssB* (Δ*tssB*) grown in LB, LB_pH6,_ and LB_NaCl_. Membranes were reprobed wih antibody anti RNA Polymerase α. In panels B, C and D, images are representative of three independent experiments.

To investigate the transcriptional pattern of the other two T6SS loci, we constructed two additional reporter strains in which *lucFF* was under the control of the promoter regions of *tssK* and *omp* encoded in locus II and locus III, respectively (Fig 1). The expression of both fusions was also higher in the exponential phase of growth (S3B and C Fig). The expression of *omp::lucFF* followed similar pattern than that of *tssB::lucFF,* whereas the expression of *tssK::lucFF* was not upregulated by iron restriction, ionic strength and osmolarity (S4 Fig).

Of all the conditions tested, the highest luciferase levels of the *tssB::lucFF* fusion were observed when Kp52145 was grown in LB buffered to pH 6 (LB_pH6_), and in LB containing 595 mM NaCl (LB_NaCl_). Therefore, we sought to determine whether Kp52145 may exhibit T6SS-mediated antibacterial activity in these conditions. Kp52145 killed *E. coli* when *Klebsiella* was grown in LB_pH6_ (Fig 3B) and LB_NaCl_ (Fig 3C), the latter condition being in which less *E. coli* was recovered. However, this was not the case when *E. coli* was co-incubated with Kp52145 *clpV* and *tssB* mutants, indicating that Kp52145-triggered *E. coli* killing is T6SS dependent (Fig 3B and 3C). We did not observe any difference in *E. coli* recovery when co-incubated with either the wild type or the *cps* mutant (S5 Fig), reinforcing the notion that *K. pneumoniae* CPS does not interfere with T6SS-dependent killing.

To further confirm that Kp52145 T6SS is active when bacteria are grown in LB_pH6_ and LB_NaCl_, we assessed the presence of Hcp in culture supernatants. The secretion of Hcp is considered an indicator of a functional T6SS [18,30]. The Hcp1 (locus tag BN49_2467, encoded in the locus I) was fused to a VSV-G tag at the C terminus and cloned into pBAD30 to give pBADHcp1. The plasmid was transferred to the wild-type strain, and the *tssB* mutant, and Hcp1 levels were assessed by immunoblotting after induction of the *ara* promoter of pBAD with arabinose. Immunoblotting experiments using anti-VSV-G antibodies showed that Hcp1 was expressed in the cytosol of both strains. However Hcp1 was only detected in the supernatants of the wild-type strain grown in LB_pH6_ and LB_NaCl_ (Fig 3D), further confirming that the T6SS is functional and secretes Hcp, a hallmark T6SS protein, when Kp52145 is grown in LB_pH6_ and LB_NaCl_.

### Regulatory networks controlling *K. pneumoniae* T6SS expression

To further characterize the regulation of Kp52145 T6SS, we sought to identify regulators controlling the expression of the T6SS when *Klebsiella* is grown under different conditions. We focused on the regulation of locus I since our previous results showed that the expression pattern of this locus correlates with the function of the T6SS using as read-outs antibacterial competition and Hcp1 secretion. The *tssB::lucFF* transcriptional fusion was introduced into *phoQ, mgrB, rcsB, h-ns, fur, hfq, rpoS, rpoN,* and *pmrAB* mutants, and the amount of light determined. When the reporter strains were grown in LB, the expression of *tssB::lucFF* was significantly higher in the *h-ns*, *mgrB* and *rcsB* mutant backgrounds than in the wild-type one (Fig 4A). In contrast, the activity of the transcriptional fusion was lower in the *phoQ, pmrAB, hfq, fur, rpoS,* and *rpoN* backgrounds than in the wild-type one (Fig 4A). As anticipated*, E. coli* was not killed when co-incubated with *rpoN, rpoS, hfq, fur* and *pmrAB* mutants (Fig 4B). However, the increased expression of the fusion in the *h-ns*, *mgrB* and *rcsB* mutant backgrounds resulted in increased activity of the T6SS because co-incubation of the *h-ns*, *mgrB* and *rcsB* mutants with *E. coli* in LB resulted in a significant reduction in *E. coli* survival (Fig 4B). The fact that studies from our laboratory have demonstrated that *phoPQ* expression is up regulated in the *mgrB* and *rcsB* mutant backgrounds [31,32] suggests that the up-regulation of *tssB::lucFF* in the *mgrB* and *rcsB* backgrounds was due to up-regulation of *phoPQ*. Supporting this hypothesis, the activity of the transcriptional fusion was downregulated in the double mutants *mgrB-phoQ* and *rcsB-phoQ* backgrounds (Fig 4A), and *E. coli* was not killed when co-incubated with the double mutants *mgrB-phoQ* and *rcsB-phoQ* (Fig 4B).

**FIGURE 4.**
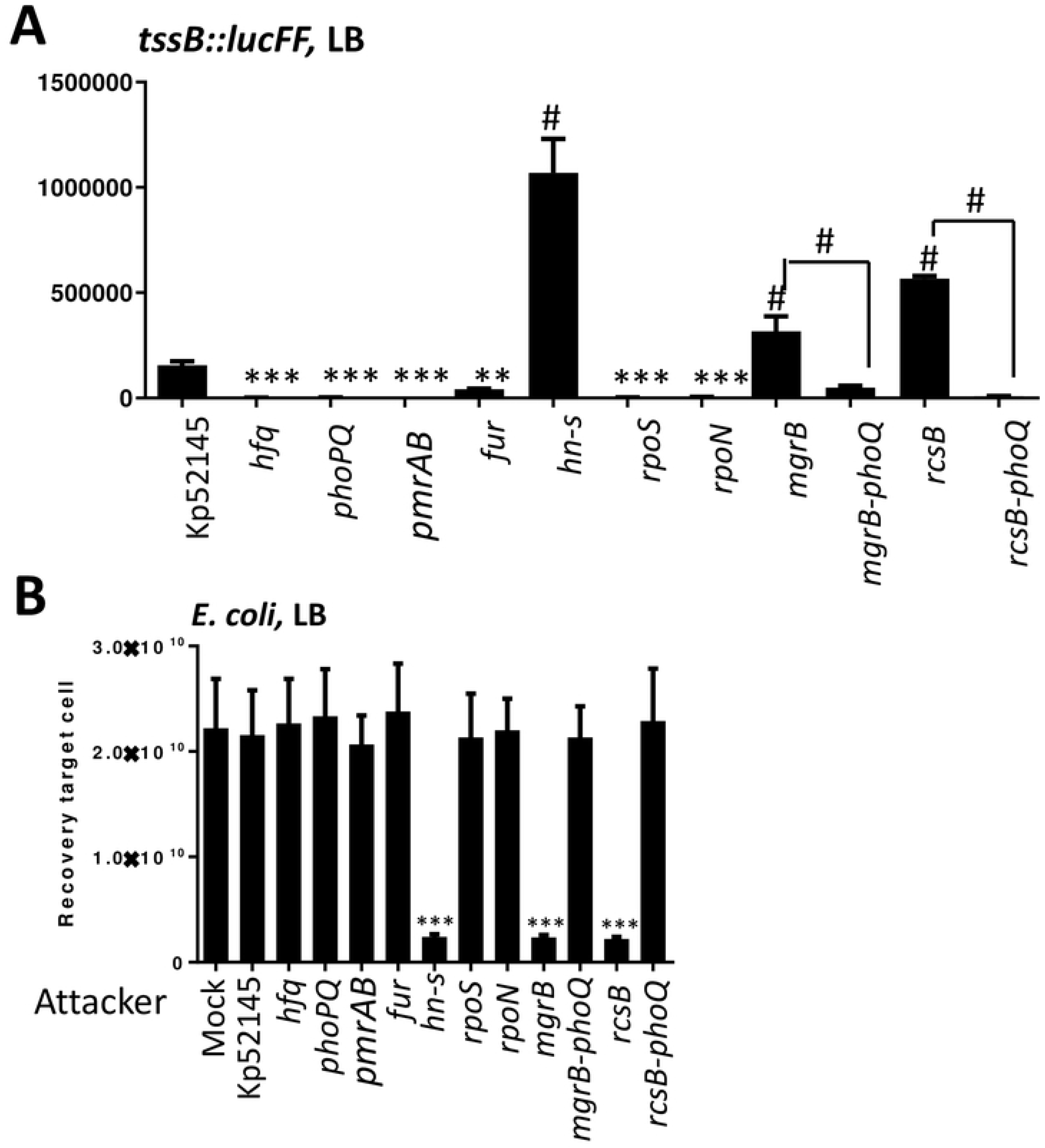
Screen of transcriptional factors controlling *K. pneumoniae* T6SS expression. (A) Analysis of the T6SS expression by *K. pneumoniae* 52.145 (Kp52145), 52145-Δ*hfq* (*hfq*), 52145-Δ*phoPQGB* (*phoPQ*), 52145-Δ*pmrAB* (*pmrAB*), 52145-Δ*fur* (*fur*), 52145-Δ*hn-s* (*hn-s*), 52145-Δ*rpoS* (*rpoS*), 52145-Δ*rpoN* (*rpoN*), 52145-Δ*mgrB* (*mgrB*), 52145-Δ*mgrB-*Δ*phoPQGB* (*mgrB-phoPQ*), 52145-Δ*rcsB (rcsB),* and 52145-Δ*rcsB-*Δ*phoPQGB* (*rcsB-phoPQ*) carrying the transcriptional fusion *tssB::lucFF* and grown in LB. Luminescence is expressed as relative light units (RLU). The data are presented as means ± the standard deviations (n = 3). *#, results are significantly different (P < 0.0001 [two-tailed t test]) from the results for Kp52145; ***, results are significantly different (P < 0.001 [two-tailed t test]) from the results for Kp52145. (B) Bacterial killing activity mediated by *K. pneumoniae* 52.145 (Kp52145), 52145-Δ*hfq* (*hfq*), 52145-Δ*phoPQGB* (*phoPQ*), 52145-Δ*pmrAB* (*pmrAB*), 52145-Δ*fur* (*fur*), 52145-Δ*hn-s* (*hn-s*), 52145-Δ*rpoS* (*rpoS*), 52145-Δ*rpoN* (*rpoN*), 52145-Δ*mgrB* (*mgrB*), 52145-Δ*mgrB-*Δ*phoPQGB* (*mgrB-phoPQ*), 52145-Δ*rcsB (rcsB),* and 52145-Δ*rcsB-*Δ*phoPQGB* (*rcsB-phoPQ*) against *E. coli* MG1655. Mock, PBS-treated *E. coli*. Number of recovered target cells following 6 h incubation in LB is indicated. The data are presented as means ± the standard deviations (n = 3). ***, results are significantly different (P < 0.001 [two-tailed t test]) from the results for Kp52145.

The expression of the fusion *tssB::lucFF* in the different strains’ backgrounds was also assessed growing the reporter strains in LB_pH6_ and LB_NaCl_. Neither in LB_pH6_ nor in LB_NaCl_ we observed the up-regulation of the fusion in any mutant background (S6A and B Fig). In contrast, the activity of the fusion was significantly down-regulated in the *phoQ, pmrAB, hfq, fur, rpoS*, and *rpoN* mutant backgrounds in both media (S6A and B Fig). As expected, the *phoQ, pmrAB, hfq, fur, rpoS*, and *rpoN* mutants did not kill *E. coli* when grown in either LB_pH6_ or LB_NaCl._ (S6C and D Fig). The activity of the *tssB::lucFF* fusion was downregulated in the *h-ns* mutant background only in LB_pH6_ (S6A Fig), and this result correlated with a lack of killing of the *E. coli* prey in LB_pH6_ (S6C Fig).

Collectively, these findings uncovered the correlation between the transcription of the T6SS and the ability to kill *E. coli*. Whereas H-NS repressed the T6SS in LB, PhoPQ, PmrAB, Hfq, Fur, RpoS and RpoN positively regulated the T6SS in all the conditions tested.

### Expression and activity of *K. pneumoniae* T6SS is enhanced by antimicrobial peptides

Polymyxin B and E (colistin) are antimicrobial peptides used as the “last-line” antibiotics against multidrug resistant *K. pneumoniae* infections. We and others have reported that challenging *Klebsiella* with polymyxins induces transcriptional changes affecting the pathogen pathophysiology [32-34]. We sought to establish whether incubation of Kp52145 with sub lethal concentrations of polymyxins will affect the activity of the T6SS. Incubation of Kp52145 with sub lethal concentrations of either polymyxin B or colistin resulted in a significant decrease of *E. coli* recovery in LB (Fig 5A). When the experiments were performed testing Kp52145 *clpV* mutant, we did not detect any decrease in the recovery of *E. coli*, demonstrating that the observed inhibition is T6SS-dependent (Fig 5A). These findings prompted us to investigate whether other antimicrobial peptides will also increase the activity of Kp52145 T6SS. We tested human β-defensin 3 (HBD3) because the levels of this peptide increase several fold during human pneumonia [35]. A sub lethal concentration of HBD3 also triggered a significant decrease in the recovery of *E. coli* when co-cultured with Kp51245 but not when the *clpV* mutant was tested (Fig 5B), indicating that the increased Kp52145-mediated killing is T6SS-dependent.

**FIGURE 5.**
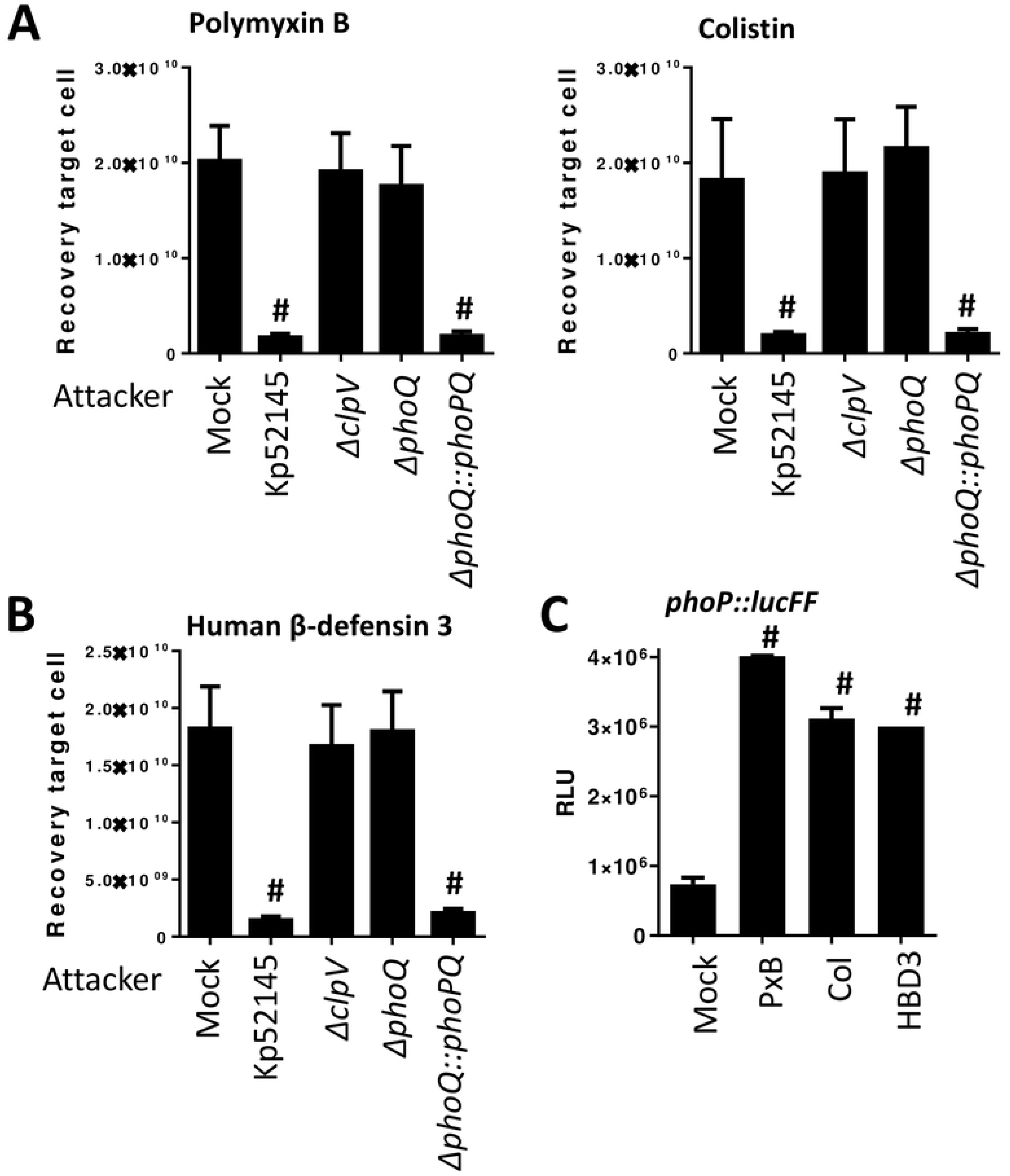
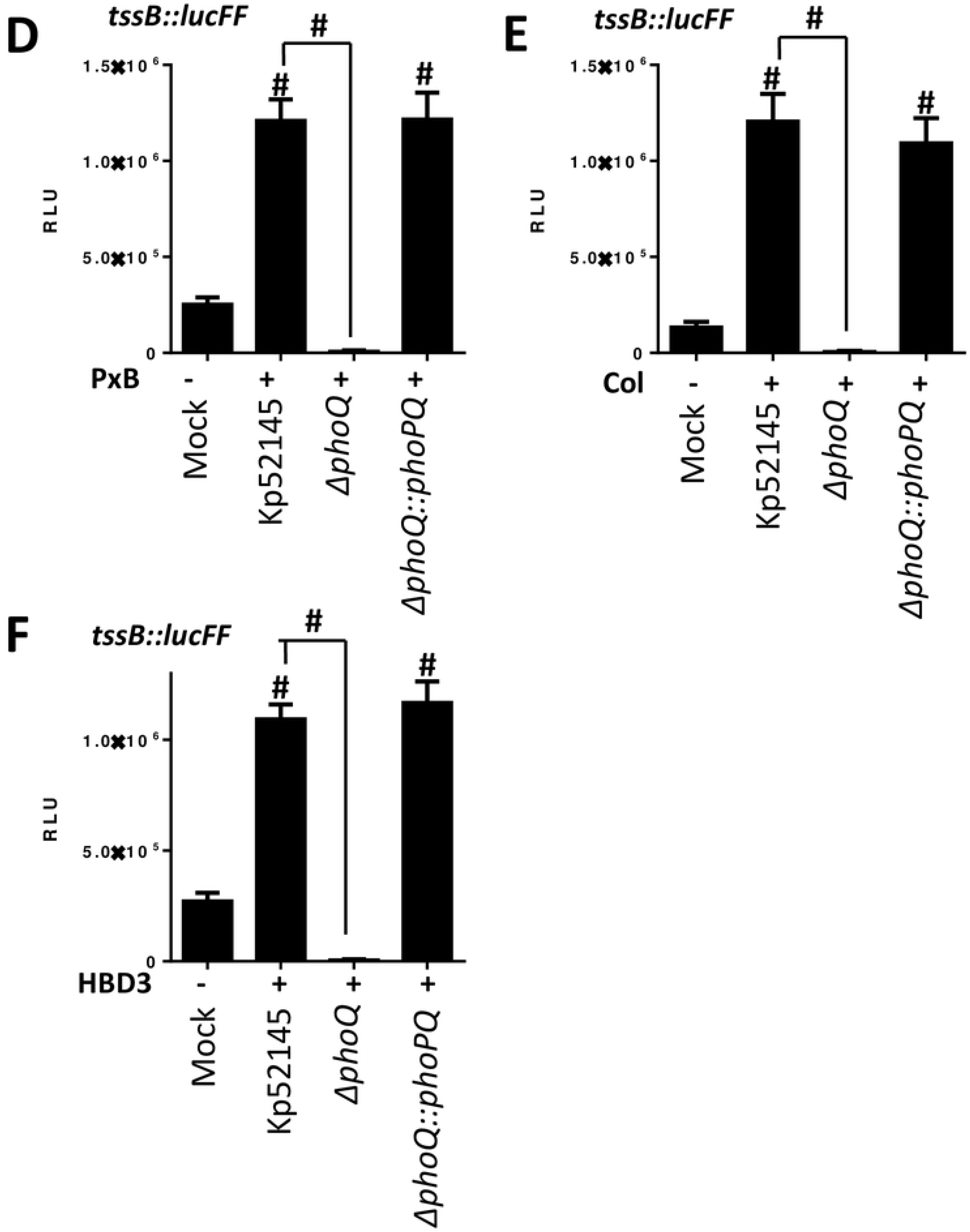
Antimicrobial peptides enhanced the expression and activity of *K. pneumoniae* T6SS. (A, B) Sub lethal concentrations of polymyxin B (0.1 μg/ml) and colistin (0.1 μg/ml), and human β-defenisn 3 (0.1 μg/ml) increase T6SS-dependent bacterial antagonism. *K. pneumoniae* CIP52.145 (Kp52145), 52145-Δ*clpV* (Δ*clpV*), 52145-Δ*phoPQGB* (Δ*phoPQ*), and 52145-Δ*phoPQGB*Com (Δ*phoPQ::phoPQ*) were incubated with the peptides for 90 min before, co-incubation with *E. coli* MG1655 prey in LB agar plates supplemented with the peptides for 6 h. Control experiments showed that incubation of *E. coli* with these sub lethal concentrations did not affect its recovery in the absence of predator bacteria. Number of recovered target cells in LB is indicated. The data are presented as means ± the standard deviations (n = 3). #, results are significantly different (P < 0.0001 [two-tailed t test]) from the results for PBS-treated (mock) *E. coli*. (C) Expression of *phoPQ* by Kp52145 carrying the transcriptional fusion *phoP::lucFF* after incubation with sub lethal concentrations of polymyxin B (PxB, 0.1 μg/ml) and colistin (Col, 0.1 μg/ml), and human β-defenisn 3 (HBD3, 0.1 μg/ml) for 90 min. Luminescence is expressed as relative light units (RLU). The data are presented as means ± the standard deviations (n = 3). #, results are significantly different (P < 0.0001 [two-tailed t test]) from the results for PBS-treated (mock) Kp52145. (D, E, F) Expression of T6SS by Kp52145, 52145-Δ*phoPQGB* (Δ*phoPQ*), and 52145-Δ*phoPQGB*Com (Δ*phoPQ::phoPQ*) carrying the transcriptional fusion *tssB::lucFF* after incubation with sub lethal concentrations of polymyxin B (PxB, 0.1 μg/ml) and colistin (Col, 0.1 μg/ml), and human β-defenisn 3 (HBD3, 0.1 μg/ml) for 90 min. Luminescence is expressed as relative light units (RLU). The data are presented as means ± the standard deviations (n = 3). #, results are significantly different (P < 0.0001 [two-tailed t test]) from the results for PBS-treated Kp52145 (mock), and the indicated comparisons.

Based on the knowledge that antimicrobial peptides activate the PhoPQ system and that PhoPQ controls the expression of Kp52145 T6SS, we asked whether antimicrobial peptides upregulate the expression of the Kp52145 T6SS in a PhoPQ-dependent manner. As we expected, sub lethal concentrations of polymyxins and HBD3 increased the activity of the *phoP::lucFF* fusion (Fig 5C). The luciferase activity of the *tssB* fusion was up regulated by polymyxins and HBD3 in a PhoPQ-dependent manner because the antimicrobial peptides did not trigger the upregulation of the transcriptional fusion in the *phoQ* mutant background (Fig 5D-F). Complementation of the *phoQ* mutant restored luciferase levels (Fig 5D-F). Altogether, these results suggest that antimicrobial peptides activate the PhoPQ two-component system which, in turn, positively regulates Kp52145 T6SS resulting in increased T6SS-dependent antimicrobial activity. Therefore, we hypothesized that antimicrobial peptides-induced T6SS-dependent killing should be abrogated in the *phoQ* mutant background. Indeed, this was the case (Fig 5A-B). Complementation of the *phoQ* mutant restored the killing potential of the *phoQ* mutant *t*o wild-type levels (Fig 5A-B).

### *K. pneumoniae* T6SS mediates intra and inter species bacterial competition in a PhoPQ-controlled manner

To establish whether Kp52145 T6SS possess the capacity to mediate intra species antagonism, we tested whether Kp52145 exerts antibacterial activity against *K. pneumoniae* strains ATCC43816, and NTUH-K2044 in a T6SS-dependent manner. Quantitative competition assays revealed that Kp52145 killed both *K. pneumoniae* strains (Fig 6A and S7A Fig). However, this was not the case when the *Klebsiella* prey strains were co-incubated with the Kp52145 *clpV* mutant (Fig 6A and S7A Fig), confirming that Kp52145-triggered killing of kin is T6SS-mediated. Similar experiments were carried to investigate whether Kp52145 T6SS mediates inter species competition. We found that Kp52145 reduced populations of *Acinetobacter baumannii* strains ATCC17978 and DSM3001 (Fig 6B and S7B Fig) and *Burkholderia cenocepacia* K56-2 (Fig 6C) in a T6SS-dependent manner. For these experiments, *A. baumannii* ATCC17978 was cured of plasmid pAB3 that suppresses the T6SS activity of this strain [36].

**FIGURE 6.**
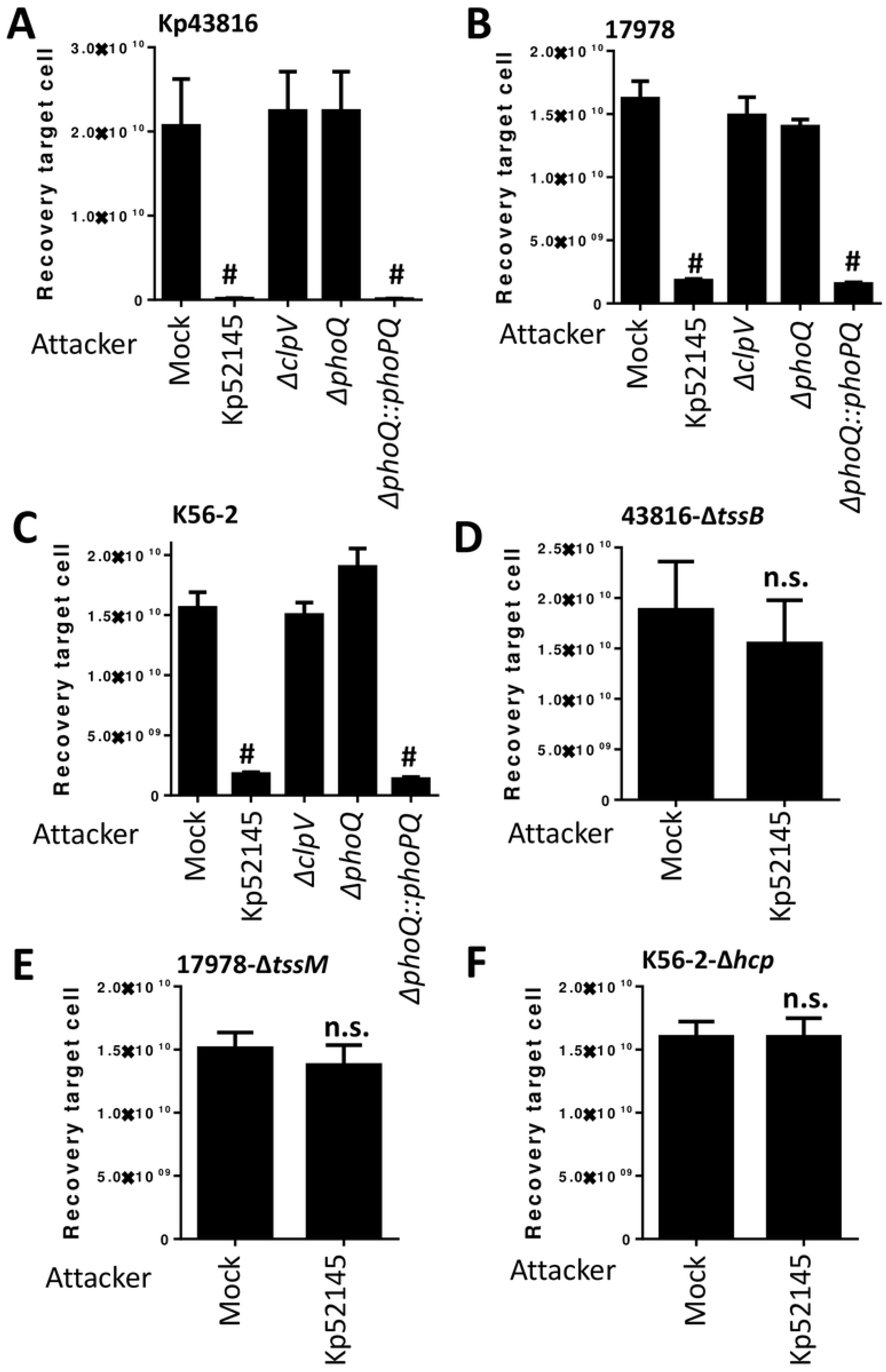
*K. pneumoniae* T6SS-dependent intra and inter species bacterial competition. (A, B, C) T6SS-dependent anti-bacterial activity as determined by recovery of target organisms *K. pneumoniae* ATCC43816 (Kp43816), *A. baumannii* ATCC17978 (17978), and *B. cenocepacia* K56-2 (K56-2) following incubation with Kp52145, 52145-Δ*clpV* (Δ*clpV*), 52145-Δ*phoPQGB* (Δ*phoPQ*), and 52145-Δ*phoPQGB*Com (Δ*phoPQ::phoPQ*). #, results are significantly different (P < 0.0001 [two-tailed t test]) from the results for PBS-treated (mock) target cell. (D, E, F) T6SS-dependent anti-bacterial activity as determined by recovery of target T6SS mutants 43816-Δ*tssB*, 17978-Δ*tssM*, and K56-2-Δ*hcp* following incubation with Kp52145. n.s., p > 0.05 (two-tailed t test) from the results for PBS-treated (mock) target cell. In all panels, the data are presented as means ± the standard deviations (n = 3).

Interestingly, Kp52145 T6SS-triggered killing of *Klebsiella*, *Acinetobacter* and *Burkholderia* was observed in LB, condition in which we have shown above Kp52145 did not exert T6SS-mediated antibacterial activity against *E. coli* prey. Previous studies in *P. aeruginosa* demonstrated that T6SS-depending killing is stimulated by T6SS activity occurring in the prey species [37]. Control experiments confirmed that *K. pneumoniae* strains ATCC43816 and NTUH-K2044, and *A. baumannii* and *B. cenocepacia* killed *E. coli* prey using their T6SS under our assay conditions (S8 Fig), uncovering the functional activity of their T6SS. When Kp52145 was co-incubated with any of the T6SS mutants of the *K. pneumoniae* strains, *A. baumannii* strains and *B. cenocepacia* there was no reduction in the recovery of the prey (Fig 6D-F and S7C-D Fig).

These results suggested that co-cultivation of *K. pneumoniae* with organisms possessing an active T6SS leads to an increased activity of the T6SS. To further investigate this hypothesis, we assessed the activity of the *tssB::lucFF* transcriptional fusion when Kp52145 was co-incubated with *K. pneumoniae* ATCC43816 and NTUH-K2044, *A. baumannii* ATCC17978 and DSM3001 and *B. cenocepacia* K56-2 wild-type and T6SS mutant strains. Co-incubation with any of the wild-type strains resulted in a significant increase in the luciferase levels (Fig 7A-C). In contrast, *tssB::lucFF* activity was not significantly different when Kp52145 was co-incubated with any of the T6SS mutants or with no bacteria (Fig 7A-C), confirming that incubation with a prey with an active T6SS results in up-regulation of *K. pneumoniae* T6SS.

**FIGURE 7.**
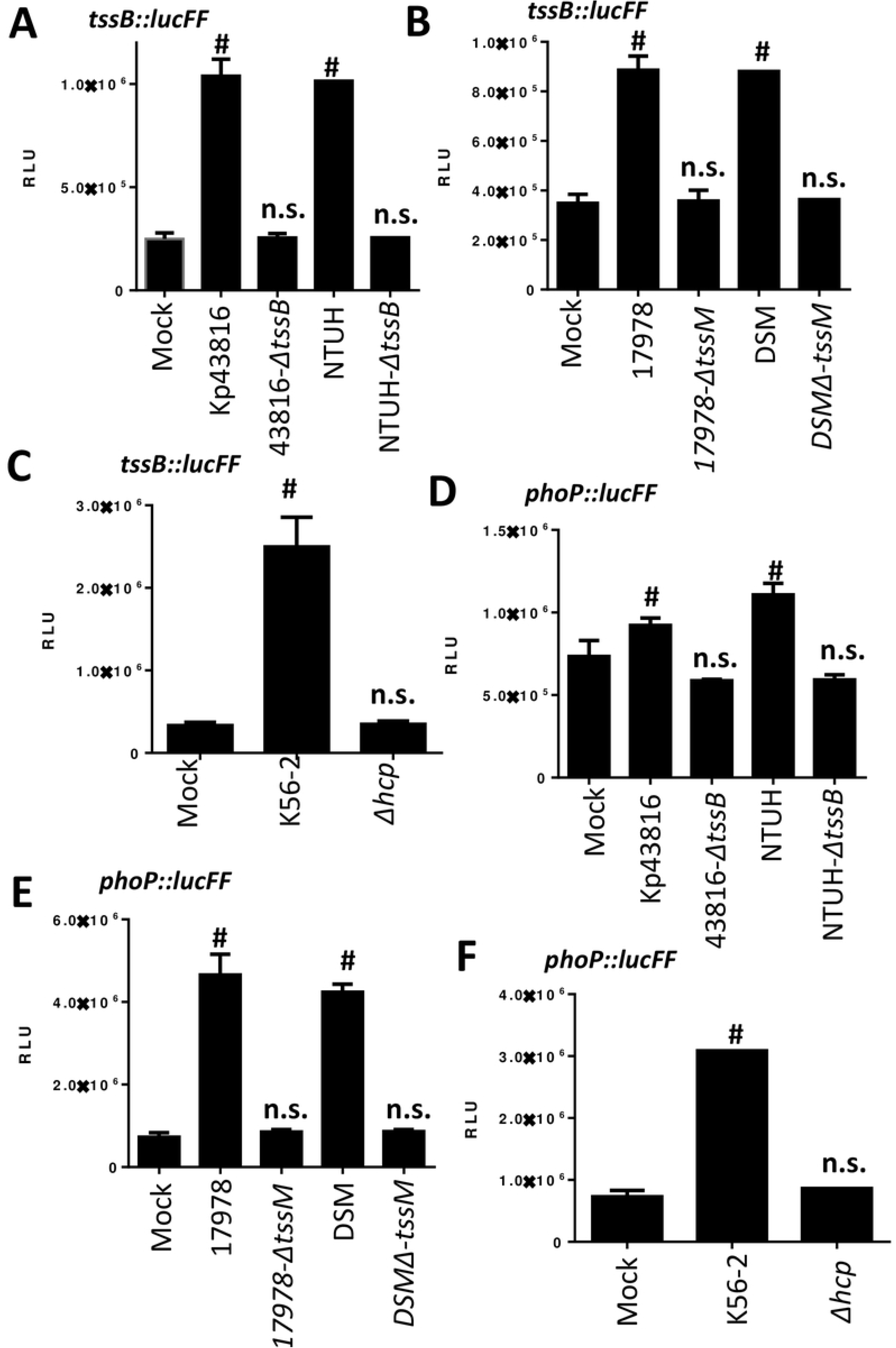
T6SS active preys activate *K. pneumoniae* T6SS and *phoPQ*. (A, B, C) Expression of T6SS by Kp52145 carrying the transcriptional fusion *tssB::lucFF* after co-incubation with *K. pneumoniae* ATCC43816 (Kp43816), 43816-Δ*tssB* (Δ*tssB*), *K. pneumoniae* NTUH-K20144 (NTUH), NTUH-Δ*tssB* (Δ*tssB*), *A. baumannii* ATCC17978 (17978), 17978-Δ*tssM, A. baumannii* DSM30011 (DSM), DSM-Δ*tssM, B. cenocepacia* K56-2 (K56-2), and K56-2-Δhcp (Δ*hcp*). (D, E, F) Expression of *phoPQ* by Kp52145 carrying the transcriptional fusion *phoP::lucFF* after co-incubation with *K. pneumoniae* ATCC43816 (Kp43816), 43816-Δ*tssB* (Δ*tssB*), *K. pneumoniae* NTUH-K20144 (NTUH), NTUH-Δ*tssB* (Δ*tssB*), A. baumannii ATCC17978 (17978), 17978-Δ*tssM, A. baumannii* DSM30011 (DSM), DSM-Δ*tssM, B. cenocepacia* K56-2 (K56-2), and K56-2-Δhcp (Δ*hcp*). In all panels, luminescence is expressed as relative light units (RLU). The data are presented as means ± the standard deviations (n = 3). #, results are significantly different (P < 0.0001 [two-tailed t test]) from the results for mock-treated Kp52145; n.s., not significant differences.

We have previously demonstrated that PhoPQ activation results in up-regulation of Kp52145 T6SS. Therefore, we sought to determine whether co-incubation with prey with an active T6SS may up-regulate *phoPQ* expression. To investigate this question, we determined the activity of the *phoP::lucFF* transcriptional fusion when Kp52145 was co-incubated with the different preys. Indeed, luciferase activity of the *phoP* fusion was up-regulated when Kp52145 was co-incubated with preys with an active T6SS (Fig 7D-F). In contrast, we observed no up-regulation of the fusion when Kp52145 was incubated with any of the T6SS mutants (Fig 7D-F). To connect PhoPQ up-regulation with that of Kp52145 T6SS, we assessed whether co-incubation with a prey with an active T6SS will up-regulate *tssB::lucFF* activity in the *phoQ* mutant background. As we anticipated, co-incubation of the *phoQ* mutant with any of the wild-type strains of *Klebsiella* (S9A Fig), *Acinetobacter* (S9B Fig) and *Burkholderia* (S9C Fig) did not result in an increase in the *tssB* activity. In fact, the luciferase levels were down regulated in the *phoQ* mutant background (S9 Fig), further confirming the role of PhoPQ to regulate T6SS expression. Complementation of the *phoQ* mutant restored luciferase levels (S9 Fig). In good agreement with these findings showing that co-incubation of the prey with the *phoQ* mutant did not upregulate the expression of *tssB::lucFF* fusion, the *phoQ* mutant did not kill any of the *Klebsiella*, *Acinetobacter* and *Burkholderia* wild-type strains (Fig 6A-C and S7A-B Fig). Complementation restored the killing ability of the *phoQ* mutant (Fig 6A-C and S7A-B Fig).

Altogether, these findings demonstrate that *K. pneumoniae* T6SS mediates intra and inter species bacterial competition. Remarkably, this antagonism of competitors is only evident when the prey possess an active T6SS. The PhoPQ two component system governs the activation of *K. pneumoniae* T6SS in bacterial competitions.

### The periplasmic domain of PhoQ is needed for the regulation of the T6SS

Studies have previously shown that the PhoQ periplasmic domain senses acid pH, divalent cations and antimicrobial peptides, and a patch of acid residues within this periplasmic sensor domain senses divalent cations and antimicrobial peptides [38]. Given that PhoPQ controls the regulation of the T6SS upon contact with prey with an active T6SS, we sought to determine whether the PhoQ periplasmic domain senses the attack of prey with an active T6SS. To address this question, the C-terminus of PhoQ was tagged with a FLAG epitope, and the construct cloned into pGP-Tn7-Cm to give pGP-Tn7-Cm*phoPQ*FLAG to complement Kp52145 *phoQ* mutant. PhoQ-FLAG tagged restored the luciferase activity of the *tssB::lucFF* transcriptional fusion upon incubation of the *phoQ* mutant with the T6SS active preys *K. pneumoniae* ATCC43817 (Fig 8A), and *A. baumannii* ATCC17978 (Fig 8B). In contrast, complementation of the *phoQ* mutant with a construct in which *phoQ* lacked the periplasmic domain, *phoQ_45-190_*, did not restore luciferase levels (Fig 8A-B). Furthermore, complementation with a *phoQ* in which the acid patch (EQGDDSE) within the periplasmic domain was mutated to (GNNNNAQ), phoQ_GNNNNAQ,_ did not result in upregulation of the *tssB::lucFF* transcriptional fusion upon incubation of the *phoQ* mutant with the T6SS active preys (Fig 8A-B). The luciferase activity of the *tssB* fusion was not up regulated by colistin when the *phoQ* mutant was complemented with either the *phoQ* lacking the periplasmic domain or the mutated acidic patch (S10A Fig). The PhoPQ-controlled up regulation of the *tssB::lucFF* transcriptional fusion when *Klebsiella* was grown in LB_pH6_ and LB_NaCl_ was not observed in any of the *phoQ* variants (S10B Fig). In good agreement with our previous results demonstrating that the PhoPQ system controls the activation of the T6SS in bacterial competitions, only the PhoQ-FLAG tagged recovered the reduction of *K. pneumoniae* ATCC43817, *A. baumannii* ATCC17978 preys upon co-incubation with *phoQ* mutant (Fig 8C-D). Complementation with *phoQ_45-190_* and phoQ_GNNNNAQ_ did not restore Kp52145 T6SS-mediated killing of the T6SS active preys (Fig 8C-D). Western blot analysis confirmed that the PhoQ FLAG tagged proteins were expressed by Kp52145 (S10C Fig).

**FIGURE 8.**
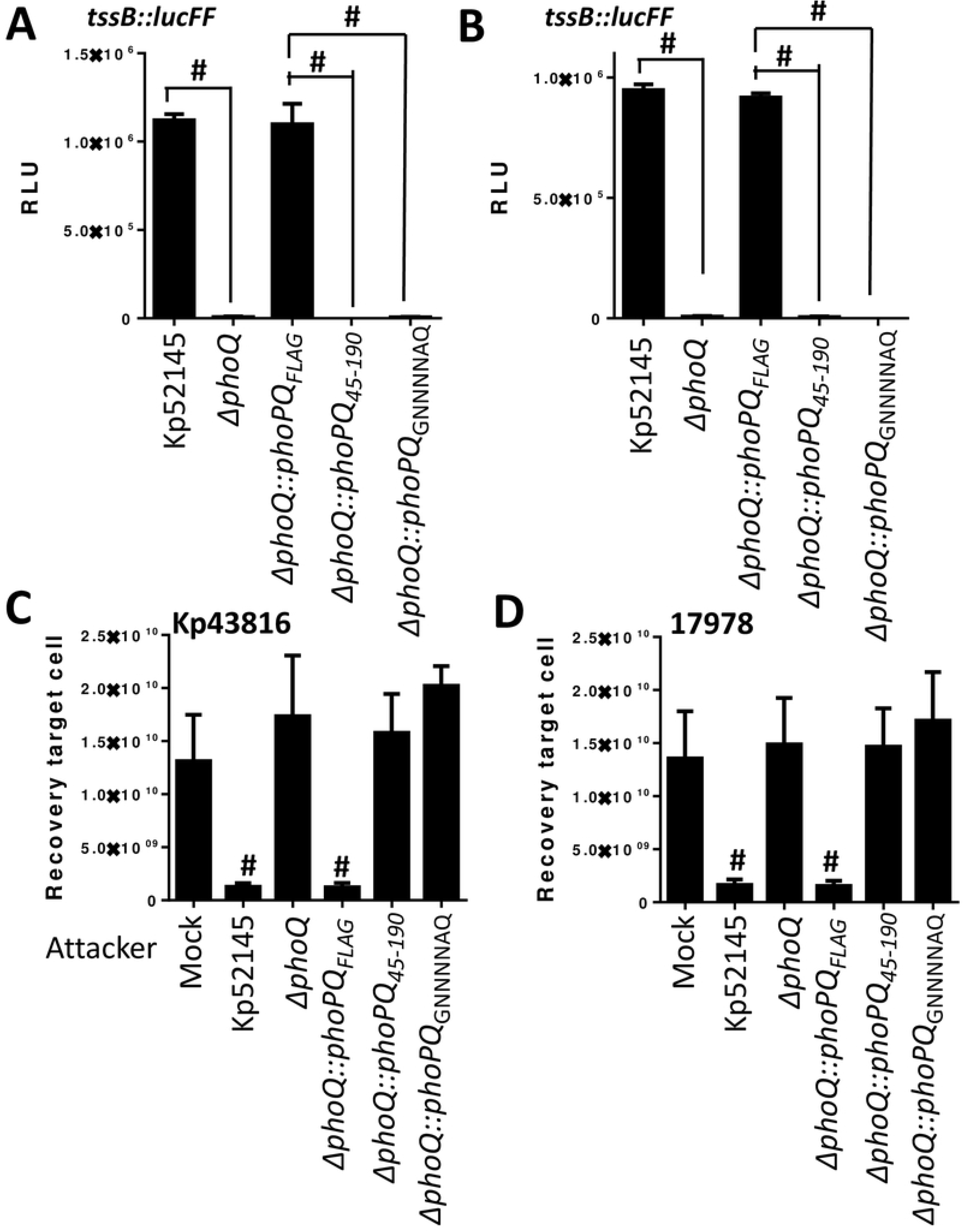
The periplasmic domain of PhoQ is needed for the regulation of the T6SS. (A, B) Expression of T6SS by Kp52145, 52145-Δ*phoPQGB* (Δ*phoPQ*), 52145-ΔphoQGBComFLAG (Δ*phoPQ::phoPQ*_FLAG_), 52145-ΔphoQGBComPhoQ_45-190_ ((Δ*phoPQ::phoPQ*_45-190_), and 52145-ΔphoQGBComPhoQ_GNNNNAQ_ ((Δ*phoPQ::phoPQ*_GNNNNAQ_) carrying the transcriptional fusion *tssB::lucFF* after incubation with *K. pneumoniae* ATCC43816 (A) and *A. baumannii* ATCC17978 (B). Luminescence is expressed as relative light units (RLU). #, results are significantly different (P < 0.0001 [two-tailed t test]) for the indicated comparisons. (C, D) T6SS-dependent anti-bacterial activity as determined by recovery of target organisms *K. pneumoniae* ATCC43816 (C), *A. baumannii* ATCC17978 (D) following incubation with Kp52145, 52145-Δ*phoPQGB* (Δ*phoPQ*), 52145-ΔphoQGBComFLAG (Δ*phoPQ::phoPQ*_FLAG_), 52145-ΔphoQGBComPhoQ_45-190_ ((Δ*phoPQ::phoPQ*_45-190_), and 52145-ΔphoQGBComPhoQ_GNNNNAQ_ ((Δ*phoPQ::phoPQ*_GNNNNAQ_). #, results are significantly different (P < 0.0001 [two-tailed t test]) from the results for mock-treated target cell. In all panels, the data are presented as means ± the standard deviations (n = 3).

Collectively, these findings revealed that the PhoQ periplasmic domain, and the acid patch within, is essential to activate *K. pneumoniae* T6SS upon incubation with T6SS active competitors, and by the last-line antibiotic colistin.

### Contribution of *K. pneumoniae* VgrGs to bacterial killing

Our bioinformatics analysis revealed that Kp52145 encodes three VgrGs, making relevant to investigate the relative contribution of each of them to bacterial killing under different conditions. We constructed single *vgrG* mutants, double mutants lacking two *vgrGs*, and a triple mutant lacking all *vgrGs*, and Kp52145-mediated T6SS killing was assessed. Quantitative killing assays using *E. coli* as a prey revealed that when *E. coli* was incubated with the *vgrG1* mutant there was no killing of the prey when strains were grown in either LB_pH6_ or LB_NaCl_ (S11A-B Fig). Complementation of the *vgrG1* mutant restored Kp52145-triggered *E. coli* killing (S11A-B Fig). Interestingly, co-incubation of *E. coli* with *vgrG2* or *vgrG4* single mutants resulted in an increase of *E. coli* recovery when bacteria were grown in LB_pH6_ and LB_NaCl_, respectively (S11A-B Fig). Complementation of the mutants restored *Klebsiella*-mediated killing of *E. coli* (S11A-B Fig). When *E. coli* was incubated with any double mutant lacking *vgrG1*, we did not observe any reduction in *E. coli* population in any growth condition (S11A-B Fig). Co-culture of *E. coli* with the double mutant *vgrG2-vgrG4* also resulted in no decrease in bacterial recovery in any growth condition (S11A-B Fig). As anticipated, the triple mutant did not kill *E. coli* in any growth condition (S11A-B Fig). VgrG1 and VgrG4 also mediated the increased activity of the T6SS upon incubation with sub lethal concentrations of colistin as observed by an increase in the recovery of *E. coli* when co-cultured with *vgrG1* and *vgrG4* single mutants (S11C Fig). Complementation of the mutants restored wild-type-triggered killing (S11C Fig). We observed no differences in the recovery of *E. coli* when co-cultured with either Kp52145 or the *vgrG2* mutant (S11C Fig). Altogether, these results suggest that VgrG1 plays an essential role in *Klebsiella*-mediated killing whereas VgrG2 and VgrG4 contribute to T6SS-dependent killing depending on the growth conditions.

We next sought to determine the relative contribution of each VgrG in inter, and intra bacterial species killing. Co-culture of the *vgrG1* mutant with *K. pneumoniae* ATCC43816 (Fig 9A), *A. baumannii* ATCC17978 (Fig 9B), and *B. cenocepacia* K56-2 (Fig 9C) resulted in no killing of any the preys. Interestingly, we observed no increase in the recovery of any of the preys when co-incubated with the *vgrG2* mutant. In contrast, co-incubation of the preys with *vrgG4* mutant resulted in an increase in the recovery of *Klebsiella* (Fig 9A), *Acinetobacter* (Fig 9B) and *Burkholderia* (Fig 9C). Complementation of *vgrG1* and *vgrG4* mutants restored Kp52145-dependent killing of the preys (Fig 9A-C), demonstrating that Kp52145 T6SS-dependent intra and inter species bacterial competition is dependent on VgrG1 and VgrG4.

**FIGURE 9.**
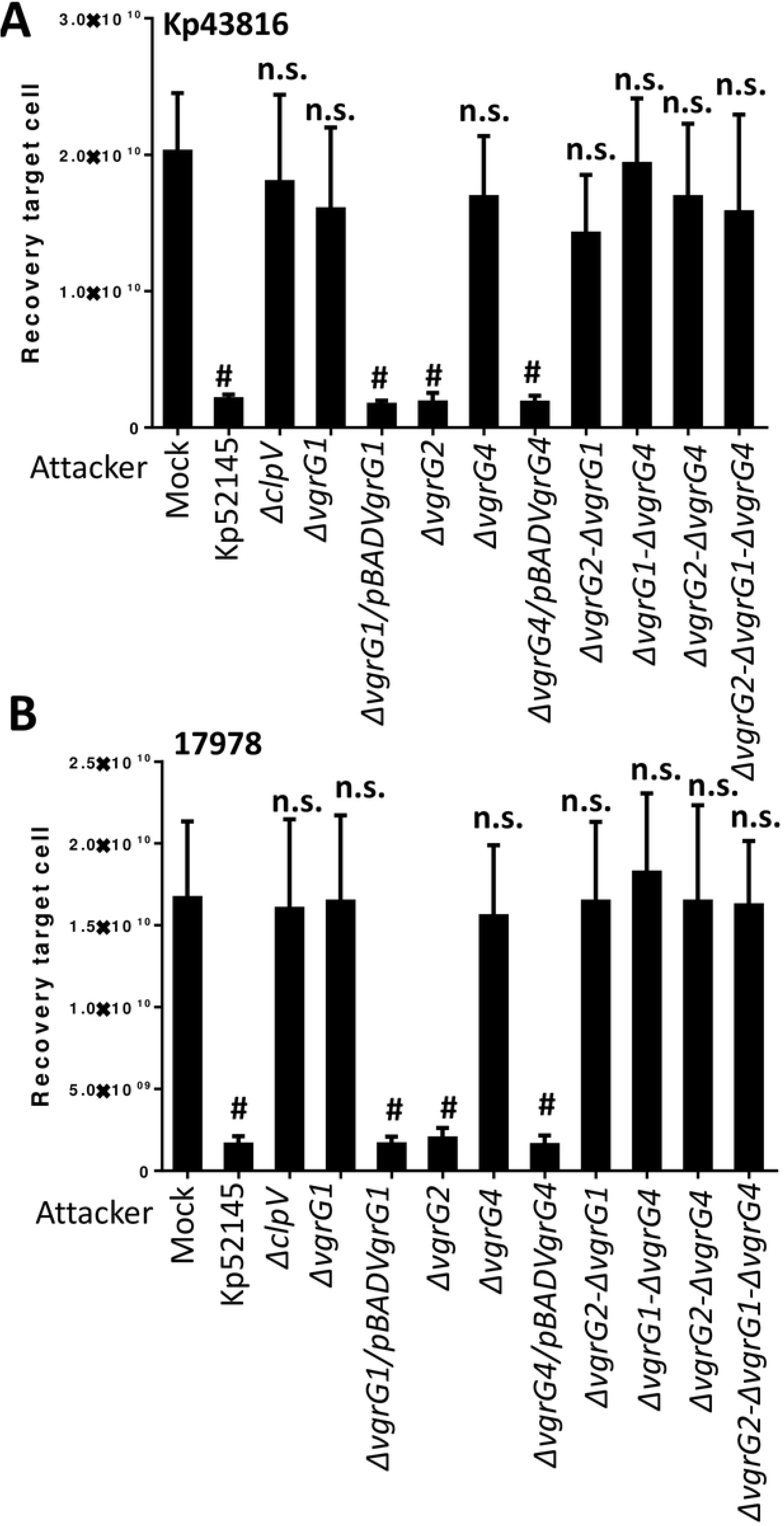
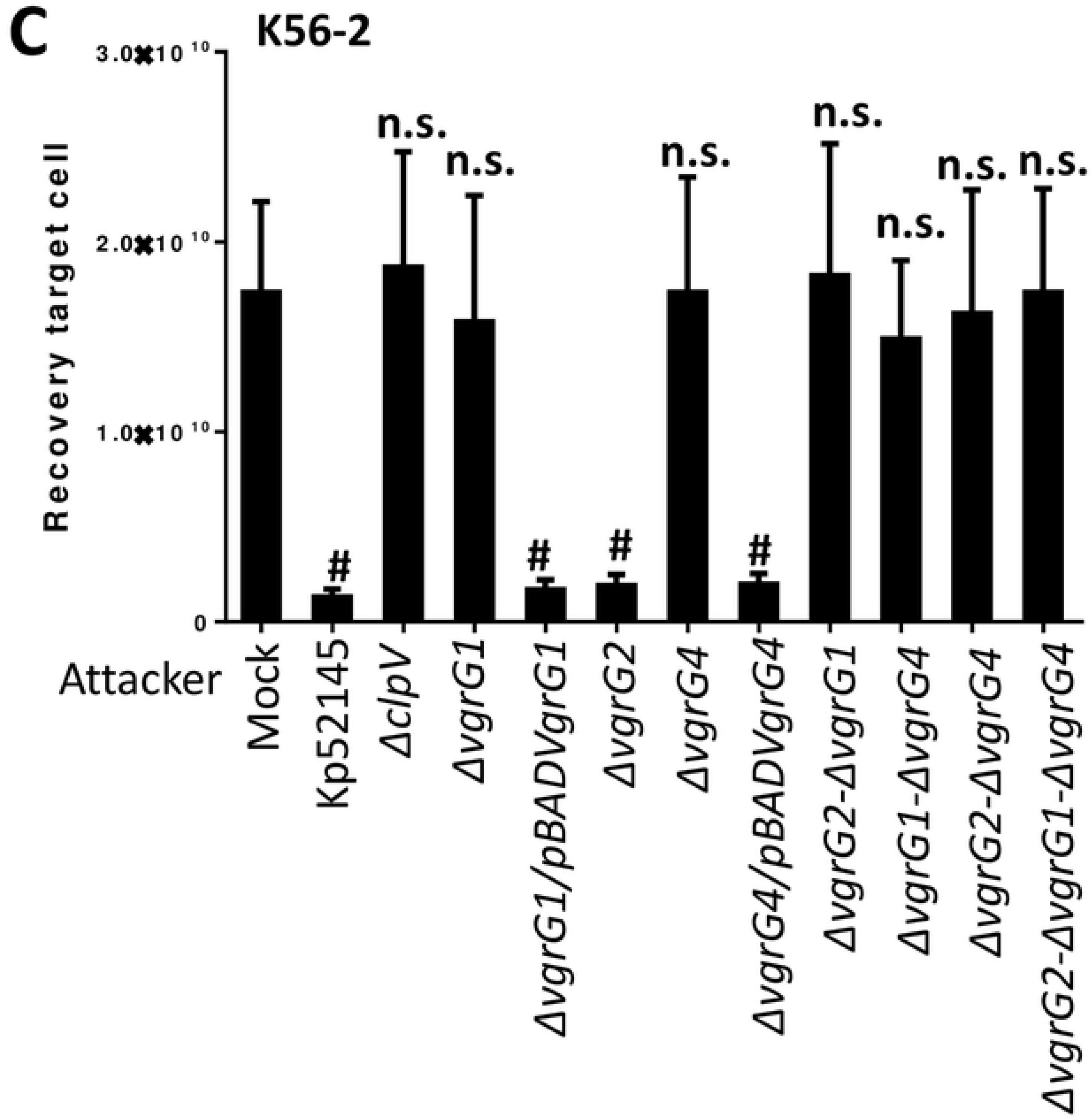
Role of *K. pneumoniae* T6SS VgrGs in bacterial competition. (A, B, C) T6SS-dependent anti-bacterial activity as determined by recovery of target organism *K. pneumoniae* ATCC43816 (Kp43816), *A. baumannii* ATCC17978 (17978), and *B. cenocepacia* K56-2 (K56-2) following incubation with Kp52145, 52145-Δ*clpV* (Δ*clpV*), 52145-Δ*vgrG1* (Δ*vgrG1*), 52145-Δ*vgrG1* harbouring pBADVgrG1 (Δ*vgrG1*/ pBADVgrG1), 52145-Δ*vgrG2* (Δ*vgrG2*), 52145-Δ*vgrG4* (Δ*vgrG4*), 52145-Δ*vgrG4* harbouring pBADVgrG4 (Δ*vgrG4*/ pBADVgrG4), 52145-Δ*vgrG2*-Δ*vgrG1* (Δ*vgrG2*-Δ*vgrG1*), 52145-Δ*vgrG1*-Δ*vgrG4* (Δ*vgrG1*-Δ*vgrG4*), 52145-Δ*vgrG2*-Δ*vgrG4* (Δ*vgrG2*-Δ*vgrG4*), 52145-Δ*vgrG2*-Δ*vgrG1-*Δ*vgrG4* (Δ*vgrG2*-Δ*vgrG1-*Δ*vgrG4*). In all panels, #, results are significantly different (P < 0.0001 [two-tailed t test]) from the results for PBS-treated (mock) target cell; n.s., not significant differences. The data are presented as means ± the standard deviations (n = 3).

### 1. *K. pneumoniae* T6SS mediates fungal competitions

To determine whether Kp52145 T6SS also mediates antifungal competition, we co-incubated Kp52145 with the clinically important fungal pathogen *Candida albicans* and the model yeast *Saccharomyces cerevisiae*. Co-culture of Kp52145 with both organisms resulted in a significant decrease in the recovery of yeast and fungal cells (Fig 10A and B). When the experiments were performed using the *Klebsiella clpV* mutant the recovery of *Saccharomyces* and *Candida* was not affected, demonstrating that the observed inhibition is T6SS dependent (Fig 10A-B).

**FIGURE 10.**
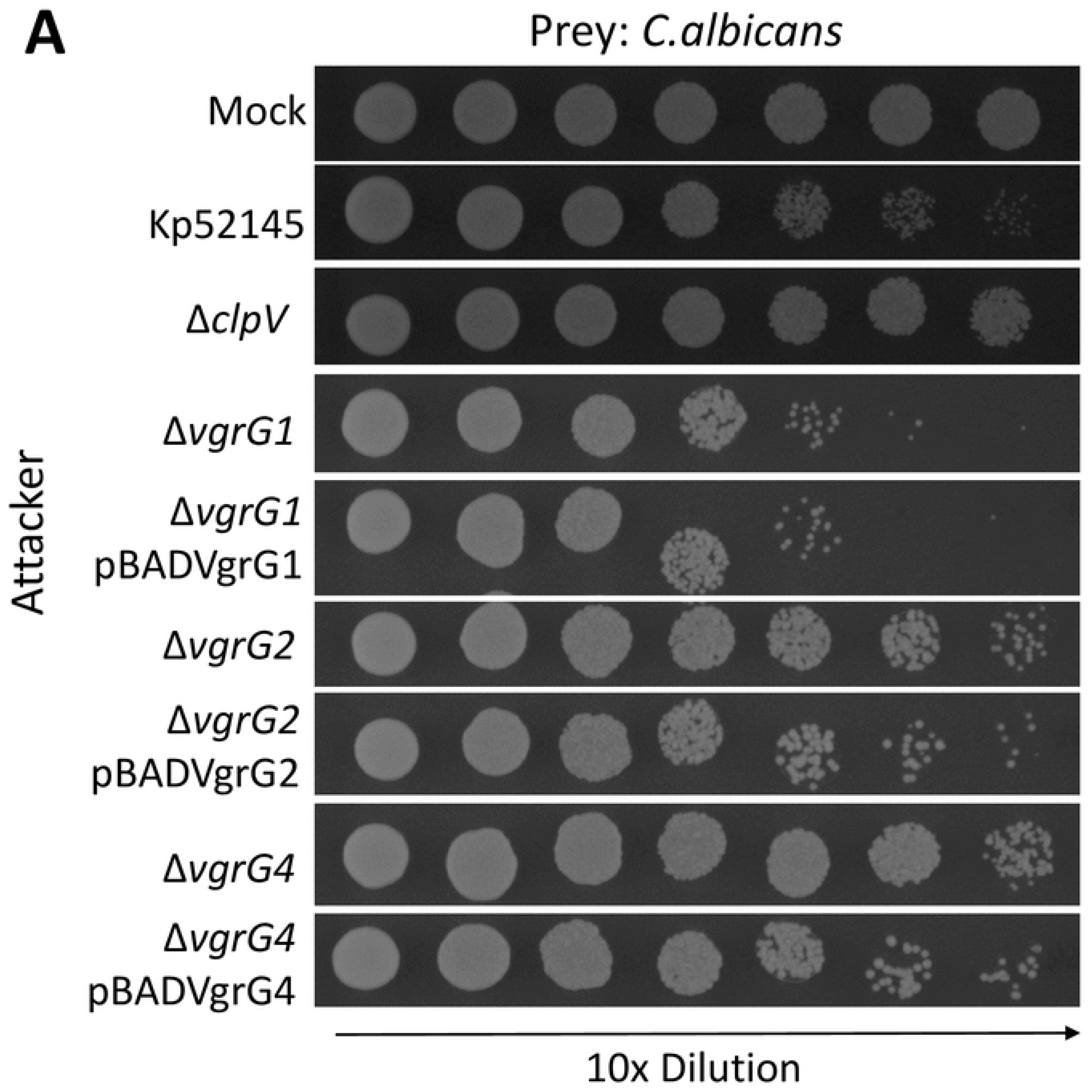
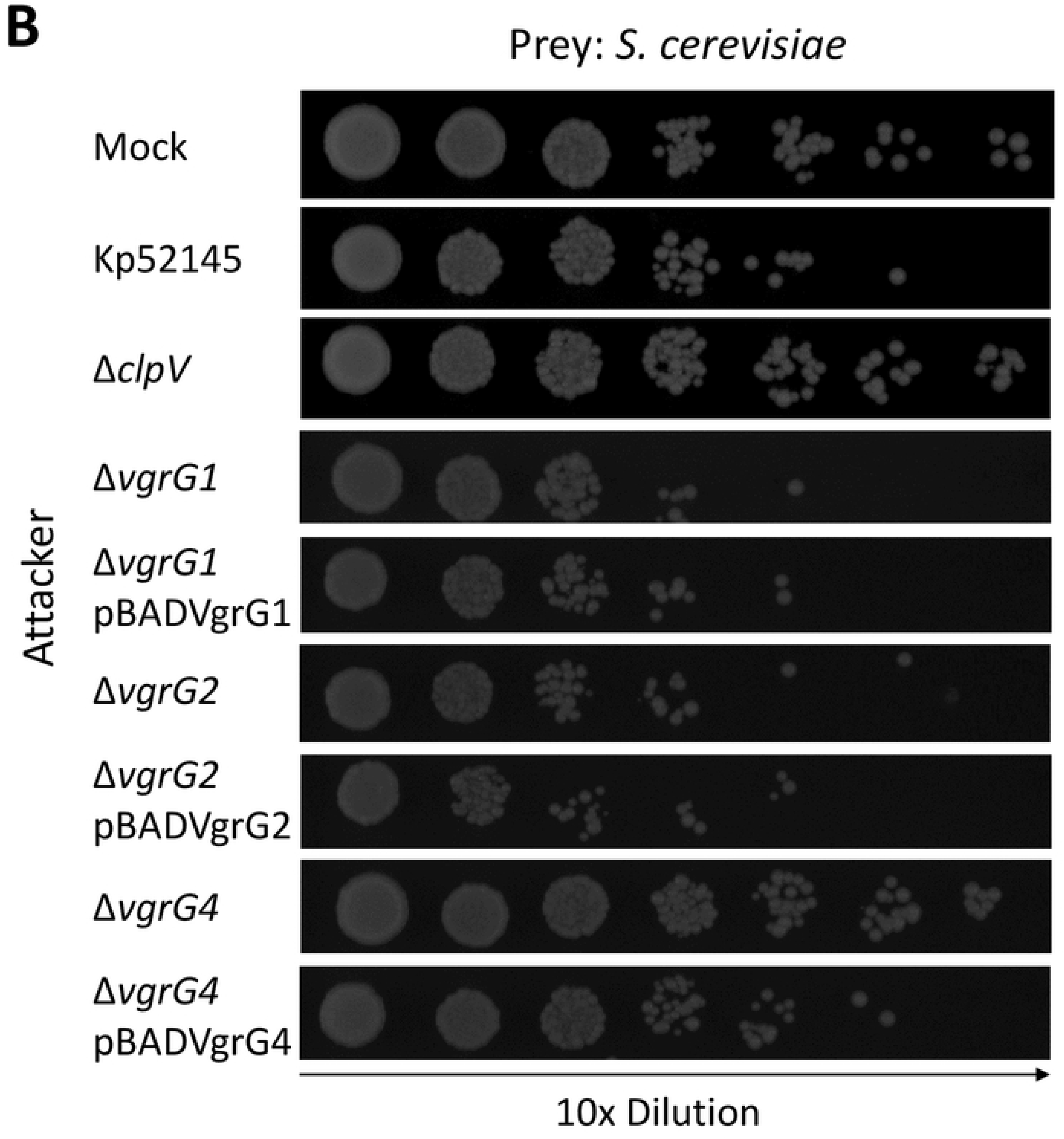
*K. pneumoniae* T6SS-dependent antifungal competition. (A, B) T6SS-dependent anti-bacterial activity as determined by recovery of target organism *C. albicans*, and *S. cerevisiae* following incubation with Kp52145, 52145-Δ*clpV* (Δ*clpV*), 52145-Δ*vgrG1* (Δ*vgrG1*), 52145-Δ*vgrG1* harbouring pBADVgrG1 (Δ*vgrG1*/ pBADVgrG1), 52145-Δ*vgrG2* (Δ*vgrG2*), 52145-Δ*vgrG2* harbouring pBADVgrG2 (Δ*vgrG2*/ pBADVgrG2), 52145-Δ*vgrG4* (Δ*vgrG4*), 52145-Δ*vgrG4* harbouring pBADVgrG4 (Δ*vgrG4*/ pBADVgrG4). After 6 h incubation in LB, target cells were recovered, diluted in PBS 0 to 10^-6^ and spotted in YNB agar plate. Mock, PBS-treated *C. albicans* and *S. cerevisiae*. Images are representative of three independent experiments.

We then investigated which of the VgrGs may play a role in Kp52145-dependent antifungal activity. Quantitative killing assays revealed an increase in *C. albicans* recovery when co-cultured with either *vgrG2* or *vgrG4* mutants (Fig 10A). Complementation of these mutants resulted in a decrease in the fungi recovered (Fig 10A), demonstrating that VgrG2 and VgrG4 are necessary for Kp52145-induced killing of *C. albicans.* VgrG4 was also necessary for Kp52145-mediated killing of *S. cerevisiae* because we observed an increase in the recovery of yeast when co-cultured with the *vgrG4* mutant (Fig 10B). In contrast, no increase in recovery was observed when *S. cerevisiae* was co-incubated with either *vgrG1* or *vgrG2* mutants. Collectively, these findings highlight a role for VgrG4 in *Klebsiella* T6SS-mediated intoxication of fungi and yeast.

### *K. pneumoniae* T6SS mutants are attenuated in the *Galleria mellonella* infection model

The *G. mellonella* infection model is well established to assess the virulence of *K. pneumoniae* [39]. Moreover, there is a good correlation between the virulence in *G. mellonella* and the mouse model [39]. To investigate whether the Kp52145 T6SS is necessary for the virulence of *K. pneumoniae* in *G. mellonella*, equal CFUs of Kp52145 and T6SS mutants were injected into *G. mellonella* and survival of the larvae was monitored over several days. Inoculation with sterile PBS into the larvae resulted in no mortality (Fig 11A). 90% of the larvae infected with 10^6^ CFUs of the *clpV* and *tssB1* mutants survived after 120 h. In contrast, only 15% of the larvae challenged with 10^6^ CFUs of Kp52145 survived (Fig 11A). Supplementary Figure 12 shows that infection with ATCC43816 and NTUH-K2044 *tssB* mutants also resulted in decrease mortality as compared to the wild-type strains, demonstrating that the contribution of the T6SS to virulence in *G. mellonella* is not dependent on the *K. pneumoniae* strain background.

**FIGURE 11.**
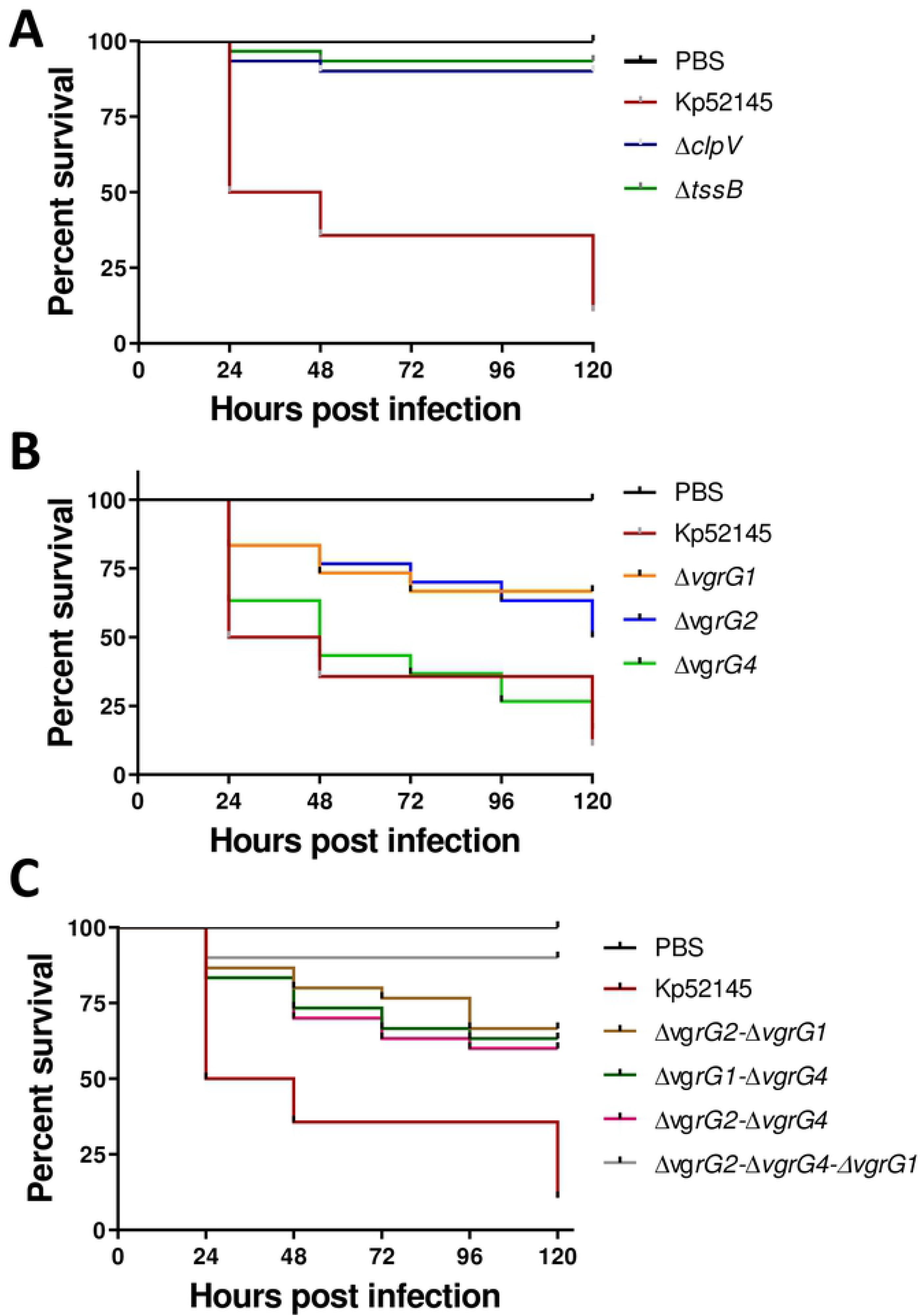
*K. pneumoniae* T6SS mutants displayed reduced virulence in the *G. mellonella* infection model. (A, B, C) Kaplan–Meier plots showing the per cent survival of *G. mellonella* over 120 h post-infection with 10^5^ organisms of the following strains: Kp52145, 52145-Δ*clpV* (Δ*clpV*), 52145-Δ*tssB* (Δ*tssB*), 52145-Δ*vgrG1* (Δ*vgrG1*), 52145-Δ*vgrG2* (Δ*vgrG2*), 52145-Δ*vgrG4* (Δ*vgrG4*), 52145-Δ*vgrG2*-Δ*vgrG1* (Δ*vgrG2*-Δ*vgrG1*), 52145-Δ*vgrG1*-Δ*vgrG4* (Δ*vgrG1*-Δ*vgrG4*), 52145-Δ*vgrG2*-Δ*vgrG4* (Δ*vgrG2*-Δ*vgrG4*), 52145-Δ*vgrG2*-Δ*vgrG1-*Δ*vgrG4* (Δ*vgrG2*-Δ*vgrG1-*Δ*vgrG4*). Thirty larvae were infected in each group. Level of significance was determined using the log-rank (Mantel–Cox) test with Bonferroni correction for multiple comparisons where applicable [a = (A) 0.0008; (B) 0.05].

We next sought to determine the relative contribution of Kp52145 VgrGs to virulence. 80% of the larvae infected with 10^6^ CFUs of the *vgrG1* and *vgr2* mutants survived whereas there was no difference in survival between larvae challenged with the *vgr4* mutant and the wild type (Fig 11B). There were no differences in the mortality triggered by any of the double *vgrG* mutants, and all of them killed only 25% of the challenged larvae after 120 h (Fig 11C). Infection with the triple mutant resulted in 100% survival of the infected larvae (Fig 11C).

Altogether, these data provide evidence demonstrating that *K. pneumoniae* T6SS is essential for virulence in the *G. mellonella* infection model. Our results also revealed that *Klebsiella* pathogenicity in *Galleria* is combinatorial with all three *vgrGs* required to result in the overall virulence phenotype.

### VgrG4 is a toxin inhibited by the antitoxin Sel1E

Bioinformatics analysis of VgrG4 revealed that it encodes a C-terminus containing a DUF2345 domain of unknown function (Fig 12A). The presence of C-terminal extensions in VgrGs is considered an indication of an effector function. Therefore, we sought to determine whether VgrG4 is an antibacterial toxin. VgrG4 was cloned into pBAD30 to control the expression of *vgrG4* by arabinose. We also cloned *vgrG1* and *vgr2* into pBAD30 to assess whether expression of any of the other *vgrGs* may have any deleterious effect in *E. coli*. The plasmids were introduced into *E. coli*, and the toxicity of the proteins upon induction assessed by plating. Figure 12B shows that expression of VgrG1 and VgrG2 in *E. coli* had no impact on *E. coli* recovery. In stark contrast, induction of VgrG4 resulted in an 80% decrease in *E. coli* recovery (Fig 12B). To assess the relative importance of VgrG4 domains for the antibacterial effect, we constructed truncated variants of VgrG4 and investigated whether they retain the antibacterial effect. Whereas VgrG4_1-517_ derivative did not affect *E. coli* recovery, VgrG4_570-899,_ containing the DUF2345 domain, was sufficient to exert an antibacterial effect (Fig 12B).

**FIGURE 12.**
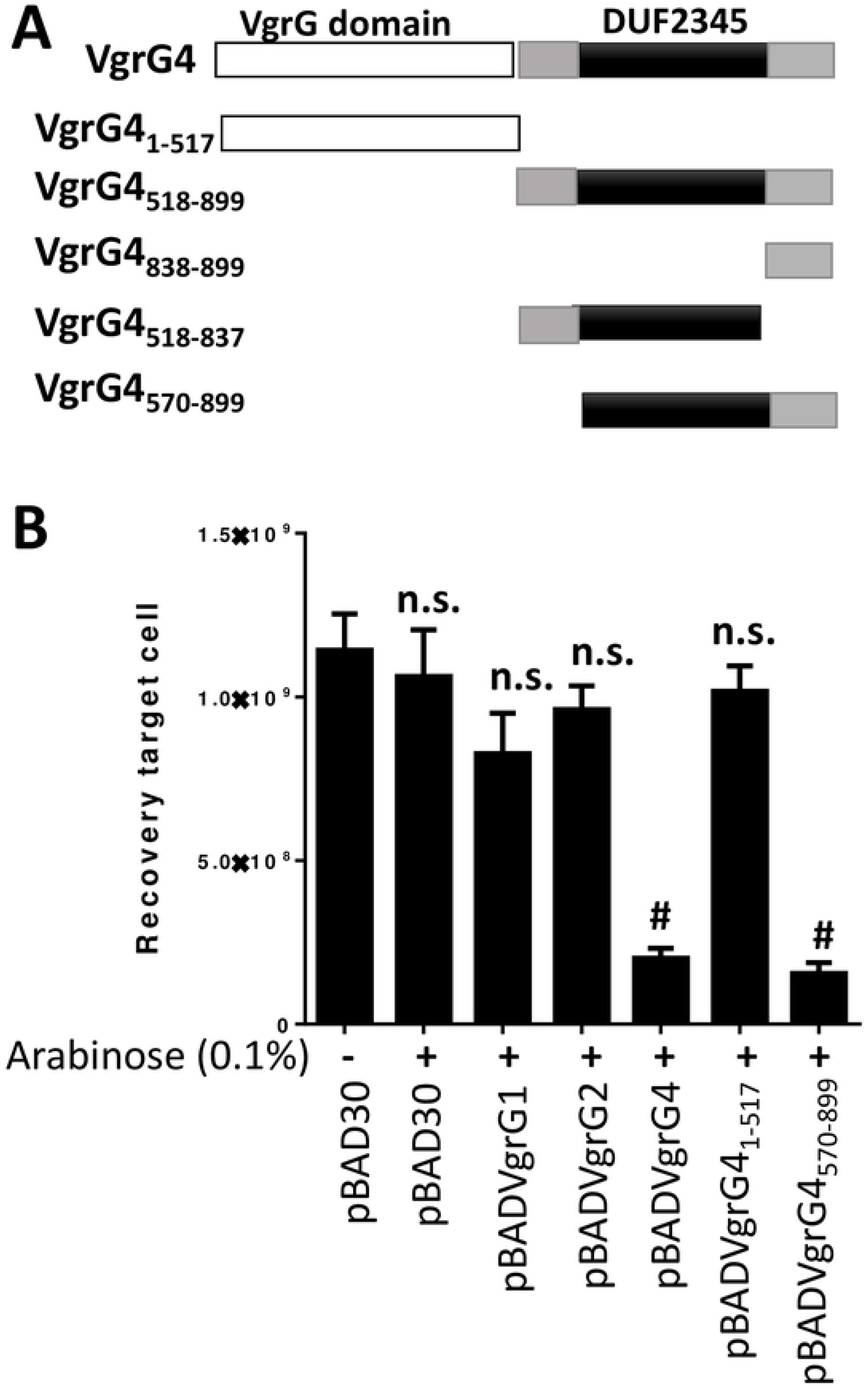
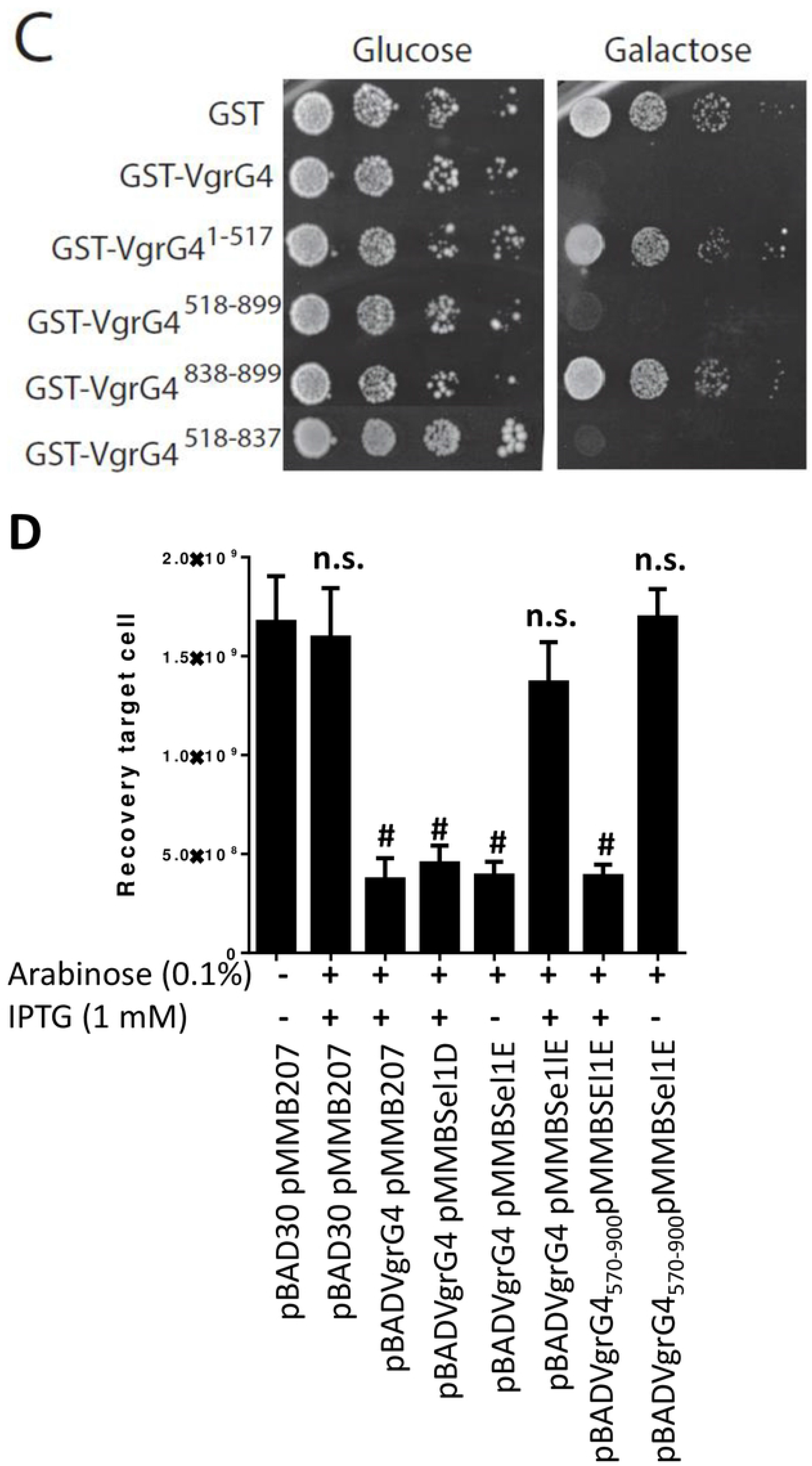
VgrG4 is a trans-kingdom effector of *K. pneumoniae* T6SS. (A) Model of VgrG4 domains. (B) Recovery of *E. coli* cells following induction of the *ara* promoter of the pBAD plasmid with arabinose (0.1%) for 60 min. #, results are significantly different (P < 0.0001 [two-tailed t test]) from the results for bacteria harbouring pBAD30 without arabinose induction; n.s., not significant differences. The data are presented as means ± the standard deviations (n = 3). (C) Growth on SD (glucose) and SG (galactose) plates of serial dilutions of wild-type YPH499 yeast cells harbouring the indicated plasmids, and therefore expressing under the control of the GAL1 promoter the corresponding proteins on galactose-based medium. Image is representative of three independent experiments. (D) Recovery of *E. coli* cells following induction of the *Ara* promoter of the pBAD plasmid with arabinose (0.1%), and the *tac* promoter of the pMMB207 plasmid with IPTG (1 mM) for 60 min. #, results are significantly different (P < 0.0001 [two-tailed t test]) from the results for bacteria harbouring pBAD30 and pMMB207 without induction; n.s., not significant differences. The data are presented as means ± the standard deviations (n = 3).

We sought to determine whether VgrG4 will also exert a toxic effect when expressing in eukaryotic cells. VgrG4, VgrG1 and VgrG2 were expressed as a GST-fusion under the control of the inducible *Gal1* promoter. We first evaluated their toxicity for the yeast cell upon induction in galactose-based medium, when the promoter is released from glucose-induced catabolic repression (S13 Fig). GST-VgrG4 caused a strong inhibition of yeast growth, whereas GST-VgrG2 from the same strain was only toxic at high incubation temperatures (37 °C) (S13 Fig). In contrast, GST-VgrG1 showed no effect on yeast growth (S13 Fig). These results revealed the broad toxin function of VgrG4. By assessing truncated variants of VgrG4, we showed that DUF2345 domain was sufficient to kill *S. cerevisiae* (Fig 12C). Control experiments confirmed the expression of the truncated VgrG4 variants in *S. cerevisiae* (S13B Fig). Collectively, these results demonstrate that VgrG4 is a bona fide T6SS toxin and that the C-terminal extension containing DUF2345 domain is sufficient for the antibacterial and anti-eukaryotic effect.

In T6SS, protection against kin cells is conferred by the production of immunity proteins that inhibit their cognate toxins. Immunity proteins are normally adjacent to the cognate toxin. Downstream of *vgr4* there are 5 genes of unknown function encoding *sel-1* repeats (Fig 1). To determine whether any of these genes assures protection against VgrG4, the genes were expressed from the pMMB207 plasmid in *E. coli* harbouring pBADVgrG4. Induction of Sel1D by IPTG did not alleviate VgrG4-triggered toxicity (Fig 12D). In contrast, induction of Sel1E abrogated VgrG4-dependent killing (Fig 12D). Furthermore, Sel1E also conferred protein against the toxicity triggered by VgrG4_570-899_ containing DUF2345 domain (Fig 12D). Altogether, these results indicate that Sel1E is the immunity protein that protects against the toxic activity of VgrG4.

### Sequence analysis and modelling of VgrG4

Sequence analyses show that the *K. pneumoniae* VgrG proteins belong to the Type VI secretion system, RhsGE-associated Vgr protein (IPR006533) protein superfamily. *P. aeruginosa* VgrG1 (hereafter PaVgrG1) belongs to the same superfamily, however, *Klebsiella* VgrG4 contains two domains that are not predicted to be found in PaVgrG1; the Putative type VI secretion system, Rhs element associated Vgr domain (IPR028244) and the Domain of unknown function DUF2345, VgrG C-terminal (IPR018769) (Fig 13A).

**FIGURE 13.**
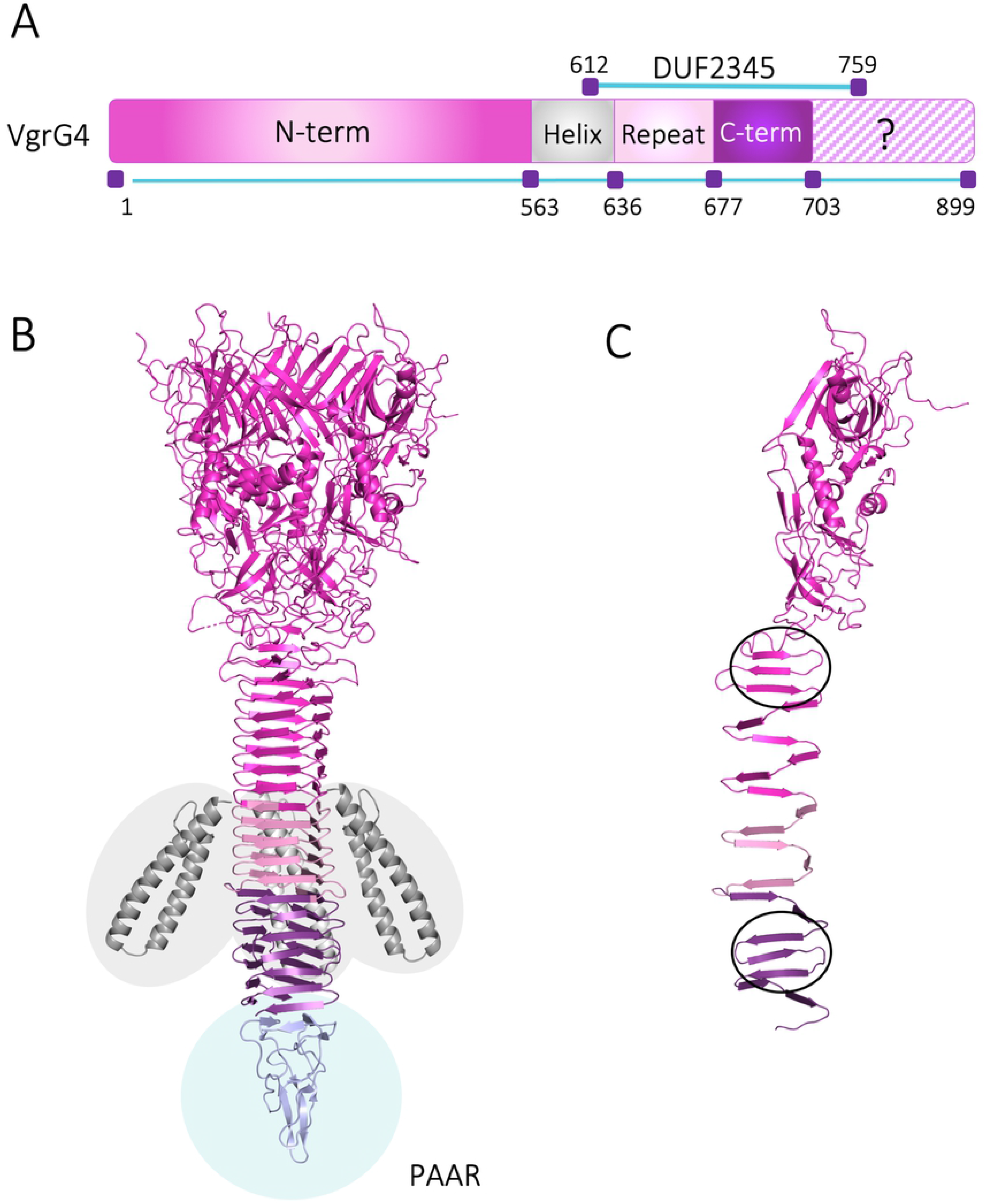
Modelling of *K. pneumoniae* VgrG4. (A) The bar represents the VgrG4 sequence and depicts the different parts that were modelled: N-term: residues 1-563, Helix: residues 564-636, Repeat: residues 637-677, C-term: residues 678-703. (B) Trimeric 3D model of VgrG4 showing the modelled regions in different colours. The separately modelled Helix part is inserted in grey in the estimated location, however, the exact orientation in relation to the rest of the protein is unknown. A PAAR protein from *V. cholerae* (PDB ID: 4jiv) is also shown to indicate the likely interaction site with the PAAR protein found in the VgrG4 locus. (C) Monomeric 3D model of VgrG4 with the two regions of antiparallel beta strands encircled.

Homology modelling is a common method used for predicting the 3D structures of proteins based on the known structure of a related protein [40] and was used to model the 3D structure of *K. pneumoniae* VgrG4. Of the two homologous PaVgrG1 structures (PDB IDs 4uhv [41], 4mtk (Sycheva et al. to be published), 4uhv was chosen as template for modelling VgrG4 since a few amino acids were missing (amino acids 1-4 and 641-643) from the 4mtk structure. The trimeric PaVgrG1 forms a hollow tube with a larger head part (N-terminus) and a narrow spike (C-terminus), connected by a neck region [41]. In the trimer structure, the spike forms a triangular-shaped beta helix consisting of 16 beta strands from each monomer. Two regions of the spike are made up of three antiparallel beta strands whereas the rest of the spike is formed by intertwined beta strands from each of the three monomers. The head contains 24 beta-strands and 4 alpha helices from each monomer as well as the irregular neck region connecting the head to the spike.

The VgrG4 protein is significantly longer (about 250 residues) than PaVgrG1 and contains additional domains not found in PaVgrG1, which made modelling more challenging. Therefore, we made separate sequence alignments for the different parts of the VgrG4 protein. VgrG4 shares a highly similar fold with the N-terminal part of PaVgrG1 and residues 1-563 (N-term) of VgrG4 was thus confidently modelled based on the PaVgrG1 crystal structure. The 3D fold of this part is almost identical to PaVgrG1 and differs only in the length of a few surface loops (Fig 13B). In the VgrG4 model, two long predicted alpha helices (Helix, residues 564-636) are located in the upper half of the spike. This region has no corresponding part in PaVgrG1 and was consequently modelled separately with I-TASSER [42], which modelled this part as two long, almost parallel, alpha helices (S14 Fig). The VgrG proteins are known to contain repeat sequences in the spike part to enable the repetitive pattern of the beta strands. For modelling of the third part of VgrG4 (Repeat), we repeated residues 551-589 in the spike part of the PaVgrG1 template to cover the VgrG residues 637-677, thereby extending the template and the part of VgrG4 that could be modelled based on it. The C-terminal part (residues 678-703) of VgrG4 was modelled based on the C-terminal end of PaVgrG1 (residues 550-645). After the end of the C-term part of the model there is a proline-rich region in the VgrG4 sequence (Fig 13A) and since prolines are unfavoured residues in beta strands (24), it is unlikely that the beta helical spike structure could continue. We were unable to model the remaining C-terminal residues (704-899) of VgrG4, and thus the folding of this region remains unknown. Figure 13C depicts the monomeric 3D modelling of VgrG4 with the two regions of beta strands circled.

### Sequence analysis and modelling of Sel1E

According to the InterPro [43] sequence analysis tool, the Sel1 proteins belong to the Tetratricopeptide-like helical domain superfamily (IPR011990) and contains Sel1-like repeats (IPR006597). The Tetratricopeptide repeat (TPR) motif consists of a 34 amino acid consensus sequence that forms two antiparallel alpha helices connected by a turn. TPR motifs are found in functionally very different protein, however, most are involved in protein-protein interactions [44]. The Sel1 repeat (SLR) sequence is very similar to the TPR motif but is usually 36-38 amino acids in length [45]. SignalP [46] and ScanProsite [47] predict that Sel1E does not contain a lipoprotein signal sequence. Localization predictions made with web servers Psortdb [48], CELLO [49] and Phobius [50] gave inconclusive results. Sel1E is about 70 residues longer than Sel1D (Fig 14A) and based on secondary structure predictions contain a long loop region at the C-terminal end with only a few predicted secondary structure elements. No information about potential domains or functions for this C-terminal region could be found through sequence analysis searches.

**FIGURE 14.**
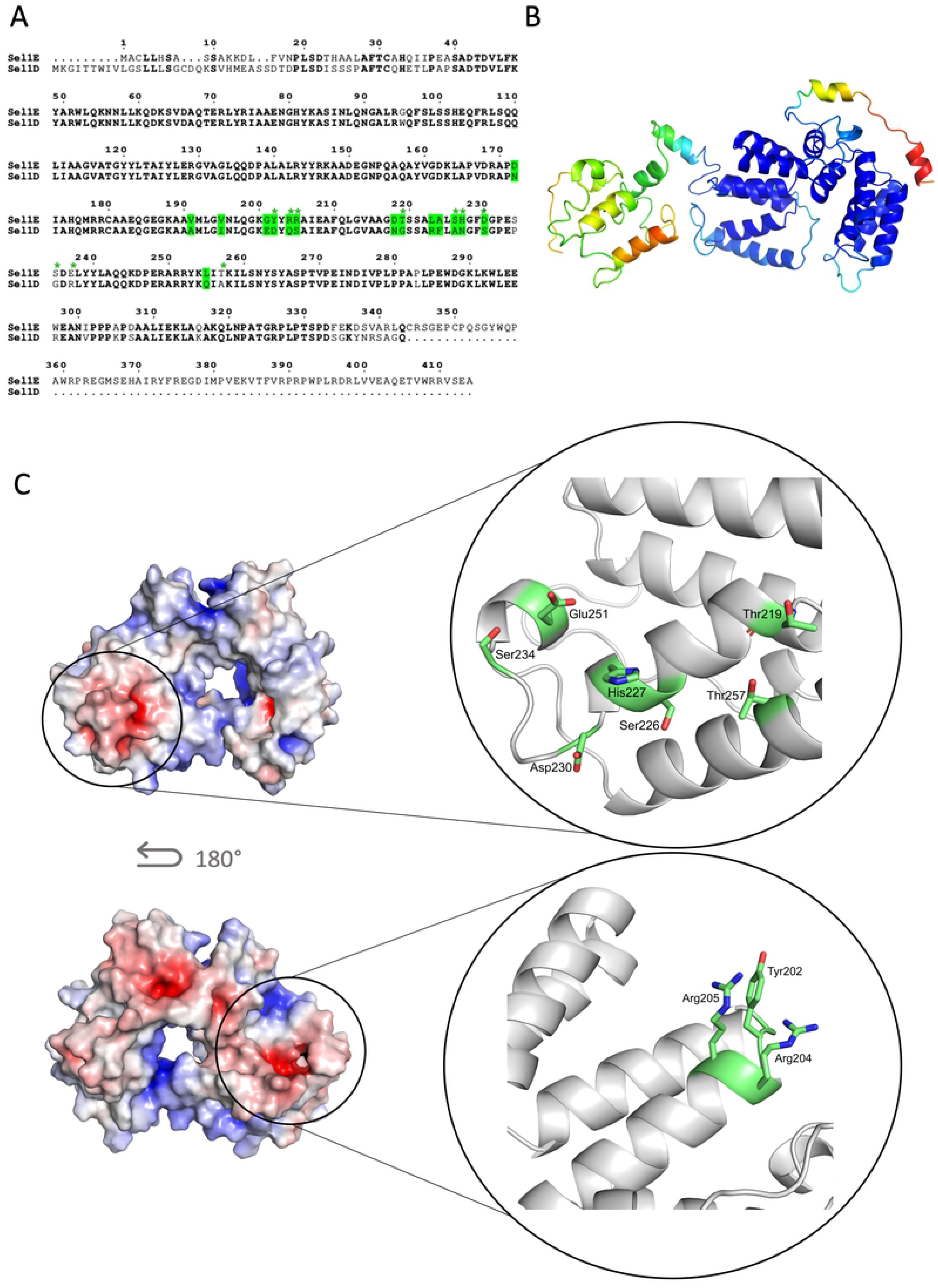
Modelling of *K. pneumoniae* Sel1E. (A) Sequence alignment between Sel1D and Sel1E showing the residues (green) that are both variable and predicted to be exposed by ConSurf. (B) MODFOLD residue accuracy prediction for the Sel1E I-TASSER model. The colour scheme for the accuracy (Å): blue (high accuracy) through green, yellow and orange to red (low accuracy). (C) Electrostatic surface potentials for Sel1E. The close-ups show the variable residues (green; marked by an asterisks in (A), which contribute to the unique electrostatic potential surface of Sel1E, see text for more details. Due to the low confidence of the C-terminal part of the model, only the region of high confidence was used for the comparison. The electrostatic surfaces were calculated with the APBS tool (Adaptive Poisson–Boltzmann Solver) in PyMOL and the colour ranges from −7 to 7.

In order to model Sel1E protein we used the threading-based method I-TASSER where we specified as a restraint that the *P. aeruginosa* Pa5087 (PDB ID: 5jkp) should be used as the structural template. The C-terminal part of the sequence not covered by the template were modelled independently by I-TASSER. Residues 28-289 were modelled with high confidence based on *P. aeruginosa* Pa5087 (5jkp). In the MODFOLD [51] residue accuracy predictions the region between residues 28-289 shows a high accuracy, whereas the C-terminal parts show lower accuracy (Fig 14A). Thus, the I-TASSER models suggest that the N-terminal end of Sel1E shares the typical fold of SLR-proteins while the structure of the C-terminal region could not be accurately determined due to a lack of homologous structures.

Next, we used the ConSurf server [52] to predict exposed residues with potential for protein-protein interactions (Fig 14A) and calculated the electrostatic potential of the protein surface for the high confident N-terminal region of the Sel1E model (Fig 14C). As our experiments show that SEl1D is not the immunity protein for VgrG4 (Fig 12D), we compared Sel1E and Sel1D and identified two regions with significant differences in the SelE and Sel1D sequences (Fig 14A, green asterisk). The first one is an acidic patch formed by Asp230 and Glu251 and several Ser and Thr residues (Fig 14C, top), whereas the second one is more positively charged due to the two Arg residues (Fig 14C, bottom). Taken together this indicates that these residues in the SLR domain of Sel1E could be important for the function of Sel1E and the interaction with VgrG4.

### VgrG4-mediated antibacterial toxicity is dependent on reactive oxygen species

T6SS antibacterial effectors can attack a number of cellular targets, including the peptidoglycan, membranes and nucleic acids. However, bioinformatics comparisons of VgrG4 with other well-characterized effectors did not provide any useful information to predict the antibacterial action/target of VgrG4. Recent evidence supports the notion that lethal insults, such as antibiotics, antimicrobial peptides and phage challenges, trigger ROS which contributes to cell killing [53-55]. We therefore speculated whether the toxic effect of VgrG4 could be mediated by ROS induction. To test this hypothesis, we asked whether VgrG4 expression up regulates the expression of *E. coli* genes that respond to ROS stress [56]. Quantitative real time PCR (RT-qPCR) experiments revealed that expression of VgrG4 induced the expression of *soxS, oxyS, oxyR, fur, and katG* (Fig 15A). The gene showing the strongest up regulation (more than 150-fold) was *soxS*, the transcriptional activator controlling the regulon that responded to ROS [56]. Moreover, VgrG4_570-899_ was sufficient to increase the expression of genes responding to ROS (Fig 15A). Interestingly, neither VgrG1 nor VgrG2 induced the expression of any of the VgrG4-upregulated genes (Fig 15A), indicating that VgrG4-induced transcriptional changes are specific. To test whether VgrG4 expression results in increased ROS levels, we used the general ROS fluorescence sensor CM-H2DCFDA [57]. We found that ROS levels were nearly 10-fold higher in *E. coli* expressing VgrG4 than in cells non-induced (S14A Fig). VgrG4_570-899_ was sufficient to increase ROS levels (S14A Fig). Single cell analysis by immunofluorescence further confirmed that VgrG4 (Fig 15B) and VgrG4_570-899_ (S14B Fig) triggered ROS in *E. coli*. Moreover, Kp52145 induced ROS in *E. coli* cells in contact with *Klebsiella* in contrast to the *vgrG4* and *clpV* mutants (S14C Fig). Complementation of *vgrG4* restored Kp52145-induction of ROS in *E. coli* (S14C Fig).

**FIGURE 15.**
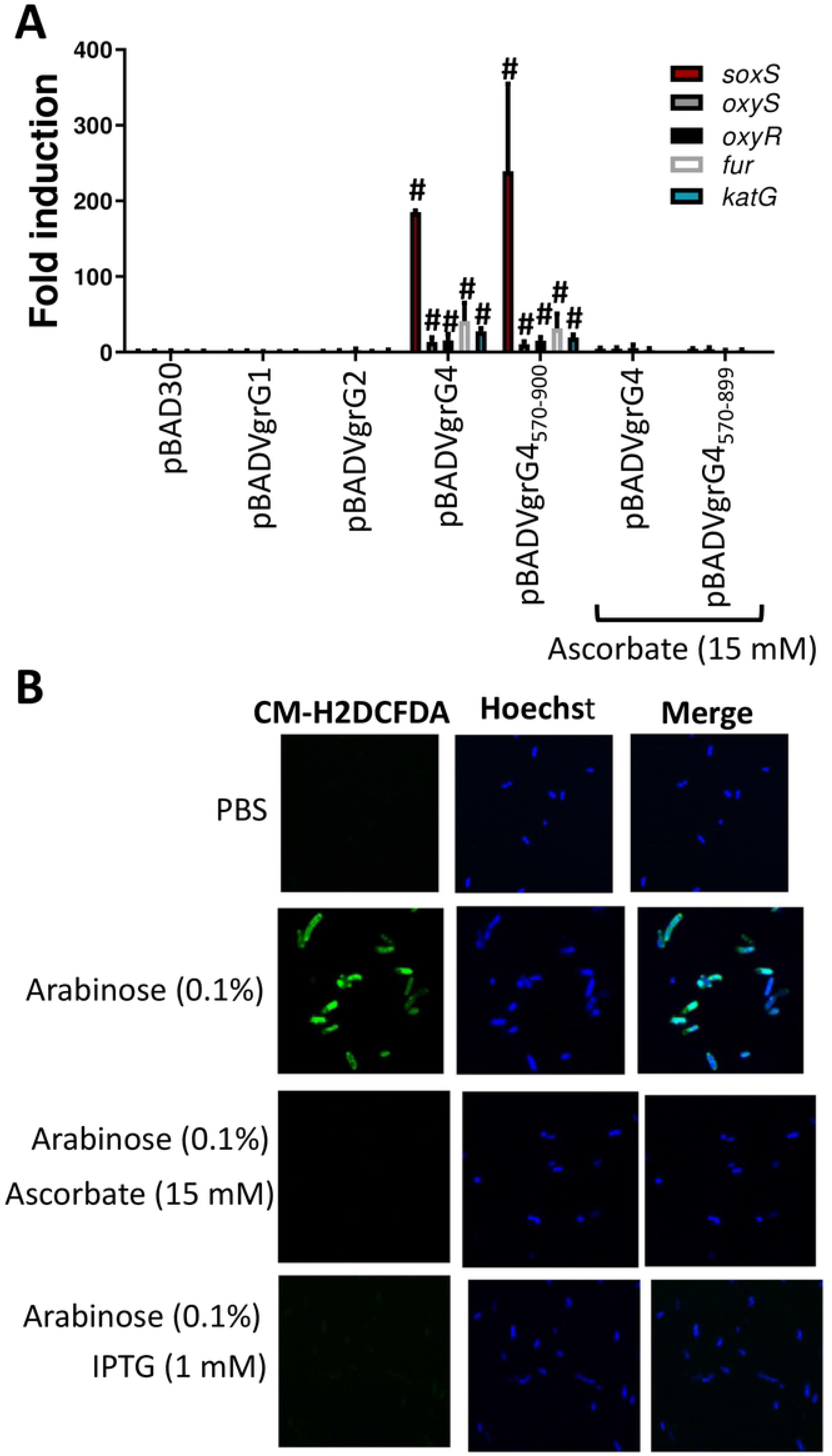
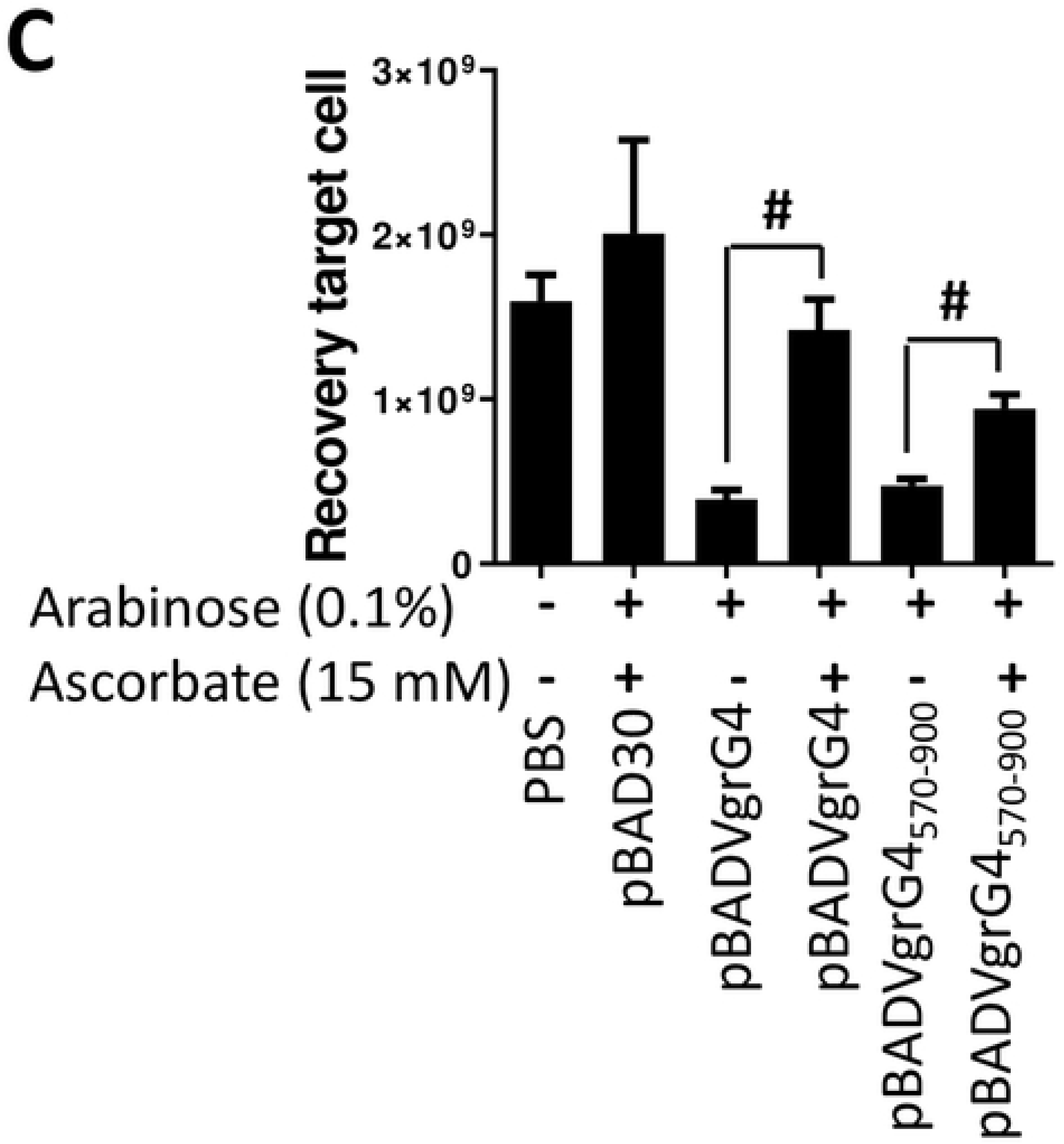
VgrG4 toxic effect is ROS-dependent. (A) *soxS, oxyS, oxyR*, *fur* and *kat* levels assessed by RT-qPCR in *E. coli* harbouring the indicated plasmids following induction of the *Ara* promoter of the pBAD plasmid with arabinose (0.1%) for 60 min. When indicated, ascorbate (15 mM) was added to supplement LB. #, results are significantly different (P < 0.0001 [two-tailed t test]) from the results for bacteria harbouring pBAD30. The data are presented as means ± the standard deviations (n = 3). (B) Single-cell analysis of ROS using the general ROS fluorescence sensor CM-H2DCFDA in *E. coli* harbouring pBADVgrG4 pMMBSel1E following induction with arabinose and IPTG for 60 min. Hoechst was used to stain bacterial DNA. Images are representative of three independent experiments. When indicated, ascorbate (15 mM) was added to supplement the culture media. (C) Recovery of *E. coli* cells following induction of the *Ara* promoter of the pBAD plasmid with arabinose (0.1%) for 60 min, and in the presence or absence of ascorbate in the growth medium. #, results are significantly different (P < 0.0001 [two-tailed t test]) for the indicated comparisons. The data are presented as means ± the standard deviations (n = 3).

To determine whether VgrG4-induced ROS mediates cell killing, we asked whether a ROS scavenger, ascorbic acid [57], may rescue *E. coli* from VgrG4-mediated killing. We found that ascorbate fully protected *E. coli* from VgrG4 and VgrG4_570-900_ -induced killing (Fig 15C). Further demonstrating the relationship between VgrG4-triggered ROS, and bacterial killing, ascorbate abolished VgrG4 and VgrG4_570-899_ -increased ROS levels (S15A Fig), and VgrG4 (Fig 15B) and VgrG4_570-899_ –induced ROS (S14B Fig) in *E. coli* cells. As expected, VgrG4 and VgrG4_570-899_ did not upregulate the expression of *E. coli* genes that respond to ROS stress when bacteria were incubated with ascorbate (Fig 15A). Collectively, these findings demonstrate that ROS generation mediates the antibacterial effects of VgrG4.

Having previously established that Sel1E protects against VgrG4-mediated toxicity, we investigated whether Sel1E prevents VgrG4 induction of ROS. Expression of Sel1E abolished VgrG4 (Fig 15B) and VgrG4_570-899_ - induced ROS (S14B Fig) in *E. coli*. Furthermore, Kp52145 did not trigger ROS in *E. coli* cells expressing Sel1E (S14C Fig). Control experiments showed that Sel1E is not a ROS scavenger because expression of Sel1E in *E. coli* did not prevent ethanol-dependent upregulation of ROS (S15A Fig). As anticipated, VgrG4 and VgrG4_570-900_ did not upregulate the expression of genes responding to ROS in *E. coli* expressing Sel1E (S15B Fig). Altogether, these results suggest that Sel1E inhibition of VgrG4-induced ROS mediates the protective effect of Sel1E.

## DISCUSSION

T6SS systems were initially studied for their interactions with eukaryotic cells, however substantial experimental data also argue for their role in microbial competition. Seminal studies probing *P. aeruginosa*, *V. cholerae*, *A. baumannii*, *S. marcescens* and *E. coli* have uncovered shared principles in terms of the structure of the secretion apparatus, and how effectors are secreted. However, these studies have also revealed significant differences between bacteria species, making necessary to investigate different bacterial models to obtain a better picture of the intricacies of the system. In this study, we provide a comprehensive characterization of *K. pneumoniae* T6SS, demonstrating its role in microbial competition and uncovering new aspects on how bacteria regulate T6SS-mediated antagonism.

Our in silico analysis revealed a significant diversity in how the T6SS is organized in *K. pneumoniae*. Although all the strains analysed encode a complete T6SS including *vgrG1*, there is diversity in the effector(s) and accompanying immunity protein. Additionally, strains may carry additional T6SS loci characterized each of them by one *vgrG* and a set of effectors and immunity proteins. The diversity found in *K. pneumoniae* T6SS reflects the considerable genomic plasticity that is contained within this species as it has been shown in population studies of this pathogen [24]. Notably, the effectors found in the analysed genomes are not similar to any other effector found in other bacteria, suggesting that investigation of these effectors will lead to the identification of new capabilities of T6SS effectors. Future structure-function studies are warranted to capture the diversity of effectors in *K. pneumoniae* pangenome.

It was somewhat unexpected to find the lack of bactericidal activity of Kp52145 against *E. coli* prey in LB. This is one of the standard assays to demonstrate the activity of any T6SS. However, our result is not unprecedented. For example, the bactericidal activity of *E. coli* EHEC is not detected in vitro [58], and that of *P. aeruginosa* H1-T6SS is only observed in the background of mutants of the negative transcriptional regulator *retS* or the Threonine Protein Phosphorylation (TPP) pathway governing the posttranslational regulation of the T6SS [30,59]. On the contrary, the T6SS of *S. marcescens* Db10 and *V. cholerae* are constitutively active [18,60]. We can only speculate why these differences exist between species, but they may reflect the adaptation to niches with different pressures in term of microbial competition. Nonetheless, it is not unreasonable to control tightly the T6SS given the energy costs to assemble such a complex structure.

Here we have shown that environmental signals such as pH, iron, oxygen tension and osmolarity control the expression of *K. pneumoniae* T6SS. Interestingly, those conditions upregulating the expression of the system, slightly acidic pH, low oxygen tension, iron restriction, and low osmolarity, are known environmental cues which may be faced by *Klebsiella* while infecting the mucosal surfaces. The relevance of these signals in other niches, particularly the environment, is largely unknown. In fact, there is a gap in the knowledge of the environment niches which might be colonized by *K. pneumoniae*. To shed further light into the regulation of *K. pneumoniae* T6SS, we screened a panel of regulators to identify those controlling the expression of the T6SS. Our results demonstrate that H-NS is a negative regulator of *K. pneumoniae* T6SS. The fact that this is a shared feature with other bacteria supports the notion that H-NS acts as a general silencer of the T6SS [58,61,62]. Our findings also reveal that RcsB and MgrB are negative regulators of the T6SS. The latter is particularly relevant because the vast majority of *K. pneumoniae* isolates resistant to the last line antibiotic colistin encode loss of function mutations of *mgrB* [63,64]. It can be then postulated that the T6SS is upregulated in these strains, resulting in increased microbial antagonism activity. Mechanistically, we have established that the increase activity of the T6SS in the *mgrB* and *rcsB* mutant backgrounds is dependent on the upregulation of the two-component system PhoPQ. In fact, our data indicate that this system acts as a positive regulator of the T6SS. PhoPQ is widely recognized as one of the master regulators governing the bacterial surface of Gram-negative bacteria, chiefly the LPS. This is also true for *K. pneumoniae*, and, for example, the dramatic lipid A remodelling observed in *mgrB* and *rcsB* mutants is PhoPQ-dependent [31,32]. This raises an interesting question whether there is a linked between the T6SS and the lipid A structure. It may well be that the assemble of the T6SS may require a certain lipid A structure, or perhaps that *Klebsiella,* as a result of an adaptation to mucosal environment, coordinates the expression of the T6SS and the LPS to interact with the immune system and competing microbes. We envision that this scenario is relevant in the context of gut colonization. Future studies are warranted to investigate these questions.

Further supporting a connection between the T6SS and the LPS, we have shown that the LPS-interacting antimicrobial peptides, polymyxin B, colistin, and β-defensin 3, upregulated the expression of the T6SS with a concomitant increase in the bactericidal activity. The facts that these agents trigger perturbations in the cell membrane, and that PhoPQ mediated this upregulation strongly suggest that an alteration of the cell membrane is the signal sensed by *Klebsiella* to control the expression of the T6SS. It is then not surprising that the PhoQ periplasmic domain, and the acid patch within it, is essential for PhoQ-mediated regulation of the T6SS because this same domain also senses cell surface perturbations caused by antimicrobial peptides [65]. Collectively, these findings support a model in which cell surface instability is a major signal controlling the T6SS. Interestingly, similar observations have been reported for *P. aeruginosa* [66,67], membrane targeting agents such as cation chelators, and polymyxin B also activate *P. aeruginosa* H1-T6SS. However, it is important to mark that in *Pseudomonas* this activation is through posttranslational control whereas in *Klebsiella* is via transcriptional control.

Our data revealed that *K. pneumoniae* also perceives T6SS attacks from bacterial competitors, resulting in retaliation against the aggressive cell. The perception of the attack involves the PhoPQ system via transcriptional regulation of the T6SS. The fact that the periplasmic domain of PhoQ is required to sense the attack strongly argues that *Klebsiella* detects the envelope perturbation after T6SS-mediated perforation. Supporting this notion, *K. pneumoniae* does not antagonize attackers in which the T6SS was mutated. Nonetheless, we cannot rigorously exclude that *Klebsiella* might detect a common structural feature of the proteins located in the T6SS apparatus upon contact.

In this study, we also show that *K. pneumoniae* T6SS mediates the intoxication of *C. albicans* and yeast. Our results follow the landmark study demonstrating that *S. marcescens* displays antifungal activity [68]. This T6SS-dependent cross-kingdom antagonism may be a widespread feature of the T6SS because it has been already found in *P. syringae* [69], and the *Serratia* T6SS effector responsible for the fungi intoxication is also encoded by *P. aeruginosa* and *V. cholerae* [68]. The antifungal activity may result beneficial for *Klebsiella* given that fungi and *Klebsiella* colonize common sites such as the gut, and the respiratory system although it is also possible that this fungicidal action may be also important in the environment.

We have uncovered that *K. pneumoniae* has a preference for different VgrGs depending the conditions. VgrG1 is essential in all conditions tested except antifungal competition, VgrG2 is required for antibacterial competition in LB_pH6_, and antifungal competition, whereas VgrG4 is needed for antibacterial competition in LB_NaCl_, and to intoxicate kin, non-kin and fungi. Based on the in silico analysis shown in Figure 1, and the accepted notion that specific effectors are absolutely dependent on particular VgrG for delivery, we can then assume that *K. pneumoniae* exploits a different repertoire of effectors depending on the condition. Taken together, it can be postulated that *K. pneumoniae* T6SS forms at least three different assemblies, each of them characterized, at least, by one VgrG protein. Future studies are warranted to characterize structurally these putative different assemblies and to define the effector(s) delivered by each of them.

Another novel finding of this work is the identification of VgrG4 as a *K. pneumoniae* T6SS effector. To date, the only T6SS effectors characterized in *K. pneumoniae* are phospholipases [22,23]. Structurally, domain DUF2345 of VgrG4 was sufficient to intoxicate bacteria and yeast. Notably, this domain is present in other VgrGs and, therefore, it can be predicted that the effector function of this domain is widespread in different bacterial species. Flaugnatti and colleagues [70], have shown that this domain present in entero-aggregative *E. coli* VgrG1 contributes to the delivery of the effector phospholipase Tle1. Our findings do not contradict this published evidence, and in fact, we predict that this is also true in the case of the *K. pneumoniae* phospholipase effector PlD1, which is encoded downstream of VgrG4 (Fig. 1A). On the contrary, our results expand the function of DUF2345 to include trans-kingdom intoxication.

VgrG4-mediated toxic effect is dependent on the induction of ROS because a known ROS scavenger rescued bacteria from VgrG4-induced lethality. Furthermore, VgrG4-induced ROS is sensed by the prey resulting in upregulation of systems aiming to alleviate ROS-imposed stress, chiefly those controlled by SoxS. Truncation experiments revealed that the C-terminus containing DUF2345 domain was sufficient to induce ROS, and the activation of *soxS*, further connecting lethality and ROS induction. In this work, we have also identified Sel1E as the VgrG4 antitoxin. Mechanistically, Sel1E inhibition of VgrG4-induced ROS mediates the protective effect of Sel1E. It is important to note that Sel1E is not a general ROS scavenger. Instead, and similarly to other T6SS antitoxins, Sel1E will interact with VgrG4 to mitigate its toxic effect. The fact that Sel1E also blunts DUF2345-induced toxicity and ROS suggests that this domain is the region of VgrG4 interacting with Sel1E. Based on our results, the SLR domain of Sel1E might mediate these interactions as the SLR repeat proteins are commonly involved in the protein-protein interactions [71].

To our knowledge, VgrG4 is the first T6SS effector for which a ROS-dependent lethality has been conclusively demonstrated. However, we believe that this can be a more widespread feature of T6SS effectors, recently it has been shown that *V. cholerae* and *A. baylyi* T6SS attacks trigger ROS induction in the *E. coli* prey [55]. Our results add further evidence to the notion that ROS generation contributes to bacteria lethality. For example, and despite the initial controversy, substantial data support that antibiotic-induced ROS contributes to their bactericidal effect in addition to their primary lethal action [54,57]. Other lethal insults such as antimicrobial peptides, and phage attack also induce ROS and trigger the upregulation of *soxS* [55]. Taken together, these results indicate that ROS generation, and the concomitant effect of ROS in gene regulation, could be considered an early event in bacterial antagonism.

## MATERIALS AND METHODS

### Bacteria and growth conditions

Bacterial strains and plasmids used in this study are shown in Table 1. Kp52145 is a clinical isolate, serotype O1:K2, belonging to the virulent CC65 clonal complex. *Burkholderia cenocepacia* and *Acinetobacter baumannii* strains have been described elsewhere and were kindly donated by Miguel Valvano, Sarah Coulthurst, Mario Feldman and Suzana Salcedo. Unless otherwise indicated, bacteria were grown in 5-ml LB medium in a 15-ml plastic tube at 37°C in a shaker (180 rpm). When appropriate media were supplemented with antibiotics at the following concentrations: ampicillin (Amp) 100 µg/ml, trimethoprim (Tmp) 100 µg/ml, tetracycline (Tet) 12.5 µg/ml, chloramphenicol (Cm) 25 µg/ml, kanamycin (Km) 50 µg/ml, carbenicillin (Cb) 50 µg/ml, apramycin (Apra) 50 µg/ml.

### Growth curve analysis

For growth analyses, 5 μl of overnight cultures were diluted in 250 μl of LB, M9 minimal medium (5× M9 minimal salts [Sigma-Aldrich] supplemented with 2% glucose, 3 mM thiamine, 2 mM MgSO4) or modified LB, and incubated at 37°C with continuous, normal shaking in a Bioscreen C™ Automated Microbial Growth Analyzer (MTX Lab Systems, Vienna, VA, USA). Optical density (OD; 600 nm) was measured and recorded every 20 min.

### Capsule polysaccharide purification and quantification

Bacteria were grown overnight in 3 ml of LB medium (37°C, 180 rpm) with viable counts determined by dilution plating. The cultures were then harvested by centrifugation, and the cell pellet resuspended in 500 μl of sterile water. Each sample was then treated with 1% 3-(N,N-dimethyltetradecylammonio)propanesulphonate (Sigma-Aldrich; in 100 mM citric acid, pH 2.0) at 50°C for 20 min. Bacterial debris was pelleted (3,220 × g, 10 min), and 250 μl of the supernatant were transferred to a clean 15-ml glass tube. The CPS was ethanol-precipitated at −20°C for 20 min and pelleted (9,447 × g, 10 min, 4°C). After removal of the supernatant, the pellet was then dried (5 min, 90°C) and resuspended in 200 μl of sterile water. CPS quantification was undertaken by determining the concentration of uronic acid in the samples, using a modified carbazole assay as previously described [72]. All samples were tested in duplicate.

### String test

A loop was used to lift a portion of a colony from a fresh LB agar plate of *K. pneumoniae*. The formation of viscous strings of >5 mm in length indicates a positive string test [73]. The string test was done on nine randomly chosen colonies from three independent plates.

### Bioinformatics mapping of the T6SS clusters

The unidentified genes within the T6SS clusters underwent a manual annotation process based on description of similarity. Pfam and COG database searches, as well as TMHMM and predictions were performed to improve the degree of annotation confidence, if necessary. Artemis, NCBI BLAST and SecRet6 database queries [74] were also used to statistically analyse unidentified proteins within the genomes of *K. pneumoniae* strains. These programs identify clusters of T6SS core components using HMMER, the amino acid sequence of the unknown proteins within the cluster are then blasted (BLASTP with a cut off of 10^5^) to putatively identify genes.

To examine T6SS diversity across the wider *K. pneumoniae* population, we downloaded all available *K. pneumoniae* genomes from the NCBI SRA (as of November 2018), and constructed a MASH distance phylogeny. From this we selected 700 genomes to represent the phylogenetic diversity of the species. We performed de novo assembly using Spades v 3.12.0 [75] and annotation using Prokka v 2.9 [76]. The component and effector database of SecRet6 was downloaded and converted into a blastable database in Abricate (https://github.com/tseemann/abricate). Using Abricate all 700 genomes were searched against the T6SS databases. A phylogenetic tree of the 700 genomes was obtained using Parsnp [77], and the gene presence-absence data visualised against the tree using Phandango [78]. To investigate the number of orthologs of *vgrG1* we used the gene sequence from genome Kp52145 to blast against the 700 genomes using blastn. All genes returning an orthologous Blast hit of over 350 consecutive nucleotides with an e-value at or close to zero were selected and investigated by blastn against the non-redundant database to confirm their identity as likely *vgrG1* orthologs.

### Marker-less mutation in *K. pneumoniae*

PCR primers used to construct the mutants were designed using the whole genome sequence of Kp52145 (GenBank Accession No. FO834906.1). The primer pairs (Table S1) were used in separate PCR reactions to amplify ∼1000-bp fragments flanking the gene to be mutated. BamHI restriction sites internal to these flanking regions were incorporated at the end of each amplicon. Purified fragments were then polymerised and amplified as a single PCR amplicon using the primers gene_UPFWD and gene_DWNRVS. These amplicons were cloned into pGEM-T Easy (Promega), and transformed into *E. coli* C600. After EcoRI digestion of the plasmids, the purified fragments were cloned into EcoRI-digested Antarctic Phosphatase (New England Biolabs)-treated pGPI-SceI-2 suicide vector [79] to generate pGPI-SceI-2Δ_gene, and transformed into *E. coli* GT115 or SY327. pGPI-SceI-2Δ_gene was thereafter transformed into the diaminopimelate (DAP) auxotrophic *E. coli* donor strain β2163 [80], and mobilised into *K. pneumoniae* by conjugation. Selection of co-integrant clones was undertaken using LB agar supplemented with Tmp at 37°C. A second crossover reaction was then performed by conjugating the pDAI-SceI-SacB plasmid [79] into a refreshed overnight culture containing three Tmp-resistant co-integrant clones. Exconjugants were selected on LB agar supplemented with Tet at 37°C. Candidate clones were checked for susceptibility to Tmp and then confirmed by PCR. Curing of the pDAI-SceI-SacB vector was performed by plating a refreshed overnight culture of one *K. pneumoniae* mutant colony onto 6% sucrose LB agar without NaCl at 30°C for 24 h. A single clone surviving the sucrose treatment was checked for susceptibility to Tet, confirmed by PCR and named 52145-Δ_gene (Table 1). The CPS levels, the mucoviscosity, and the growth of the mutant strains were assessed as previously described. No differences were found between any of the mutants constructed and the corresponding isogenic wild-type strain.

### Complementation of *vgrG* mutants

To complement the single *vgrG* mutants, a PCR fragment comprising the coding region of each *vgrG* was amplified using primers indicated in Table S1. The amplicon was digested with EcoRI and HindII in the case of *vgrG1* and *vgrG4*, and ScaI and XbaI in the case of *vgrG2*. The fragments were gel-purified and cloned EcoRI-HindIII and ScaI-XbaI-digested pBAD30, respectively, to obtain pBADVgrG1, pBADVgrG2 and pBADVgrG4 (Table 1). These plasmid were transformed into *E.coli* β2163, and then mobilized into each of the *Klebsiella vgrG* mutants by conjugation.

### Generation of the *lucFF* reporter fusion *K. pneumoniae* strains

A 1 kpb amplicon comprising the *tssB, tssK* and *omp* gene promoter regions were amplified using Phusion® High-Fidelity DNA Polymerase and the primers protssB / protssK / proomp_FWD and protssB / protssK / proomp_RVS (Table S1). The amplicons were digested with EcoRI, gel-purified, cloned into an EcoRI-SmaI-digested, Antarctic Phosphatase (New England Biolabs) pGPL01 suicide vector and then transformed into *E. coli* GT115 cells to obtain pGPLKpnProtssB, pGPLKpnProtssK, and pGPLKpnProomp. Correct insertion of the amplicons were verified by restriction digestions with EcoRI and SmaI. Vectors were introduced into *E. coli* β2163, and then mobilised into *K. pneumoniae* strains by conjugation. Cultures were then serially diluted and checked for Amp resistance by plating on LB Cb agar at 37°C. Correct insertion of the vectors into the chromosome was confirmed by PCR using the relevant lucFF_check and promoter sequence primers (Table S1).

### Antibacterial competition assay

Attacker *K. pneumoniae* strains and preys were grown until mid-exponential phase, collected by centrifugation, washed three times with PBS, and resuspended in PBS to an OD_600_ of 1. Attacker and preys were mixed at 10:1 ratio, and the mixture spot it in a pre-warmed agar plate at 37°C. The mixture was incubated for 6 hours. Recovered cells were plated out on antibiotic selective media and viable cells were reported as recovery target cell representing the CFU per ml of the recovered prey after co-culture. *Burkholderia* strains were selected by plating on Tmp (50 μg/ml), *Acinetobacter* strains by plating on Cm (10 μg/ml), and *Klebsiella* strains by plating on Cb (50 μg/ml), All experiments were carried out with triplicate samples on at least three independent occasions.

### Antifungal competition assay

*S. cerevisiae* and *C. albicans* were grown in YPD (Peptone (20% w/v), Yeast Extract (10% w/v), Dextrose (20% w/v)) overnight at 30 °C with agitation (180 r.p.m.). Cells were then adjusted in PBS to yield 5×10^7^ cells/ml. K. *pneumoniae* strains were grown to mid exponential phase, washed three times in PBS, and the suspension adjusted to an OD_600_ of 1 in PBS. *Klebsiella* and eukaryotic cells were mixed at a 10:1 ration (predator:prey), and 50 µl of the mixture were spotted in pre-warm LB plate and incubated for 6 h at 37°C. Cells were recovered, serially diluted in PBS and plated on YNB (YNB 0.67% w/v, Dextrose 20% w/v) medium containing Km. All experiments were carried out with duplicate samples on at least three independent occasions.

### Luciferase activity

Overnight cultures of the *K. pneumoniae* reporter strains were refreshed until mid-exponential phase in the appropriated media. The cells were then pelleted, washed once in sterile PB buffer (1M disodium hydrogen orthophosphate, 1.5 mM potassium dihydrogen orthophosphate, pH 7.0) and adjusted to an OD_600_ of 1.0. One hundred microlitres of each suspension were added to an equal volume of luciferase assay reagent (1 mM d-luciferin [Synchem] in 100 mM sodium citrate buffer pH 5.0), vortexed for 5 s and then immediately measured for luminescence (expressed as relative light units [RLU]) using a GloMax 20/20 Luminometer (Promega). All strains were tested in quadruplicate from three independent cultures.

To determine luciferase activity of bacteria grown in solid media, reporter strains were incubated with preys as described in the antibacterial completion assay for 6 hours. The mixture was then recovered with 1 ml of PB buffer in a microcentrifuge tube. Bacteria were collected by centrifugation (13,300 rpm, 2 mins), washed with 1 ml of PB buffer, and adjusted to an OD_600_ of 1.0. One hundred microlitres were used to assess luminescence. All strains were tested in quadruplicate from three independent cultures.

### Site-directed mutagenesis of *phoQ*

The *phoPQ* promoter and coding region including a FLAG-tag was amplified by PCR using primers Kp52_PhoPQ_compF1 and Prom_PhoPQ_FLAG_R1, gel-purified and cloned in to pGEM-T Easy to give pGEMTPhoQPFLAG. To delete the periplasmic region of PhoQ, 45-190 aminoacids [38], we followed the protocol described by Byrappa and co-workers [81] using primers Kp52_PhoPQ_SDM_F2 and Kp52_PhoPQ_SDM_R2 (Table S1), and Vent polymerase to obtain pGEMTPhoQ_45-190_. To mutate the acidic patch, aminoacids 148 to 154, within the periplasmic region of *phoQ* from EQGDDSE to GNNNNAQ, we used primers Kp52_PhoQ_Acidic_F2 and Kp52_PhoQ_Acidic_R2 to obtain pGEMTphoQ_GNNNNAQ_. The region corresponding to *phoQ* in pGEMTPhoQ_45-190_ and pGEMTphoQ_GNNNNAQ_ were sequenced to ensure that no additional changes were introduced in the coding region. The wild-type and mutated alleles were obtained by EcoRI digestion, gel-purified and cloned into pUC18R6KT7mini-Cm vector to obtain pUC18r6KT7mini-Cm PhoQ_45-190_ and pUC18r6KT7mini-Cm phoQ_GNNNNAQ_ which were then transformed into *E. coli* C600 and thereafter into *E. coli* β2163. In addition, the transposase-containing pSTNSK-Tmp plasmid [82] was introduced to the 52145-Δ*phoQ* strain by electroporation to give 52145-Δ*phoQ* / pSTNSK-Tmp. This strain was then conjugated overnight with *E. coli* β2163 containing the *phoQ* alleles on LB agar supplemented with DAP at 30°C. The retrieved mixture was then serially diluted in sterile PBS, plated onto LB Cm agar and incubated at 42°C for 6.5 hours followed by 37°C overnight. The colonies grown were thereafter screened for resistance to Amp and susceptibility to Cm. Correct integration of the Tn7 transposon was confirmed by PCR using the primers KpnglmSup/Ptn7L and KpnglmSdown/Ptn7R as previously described [31,83]. Additionally, presence of the *phoQ* gene was PCR confirmed using Kp52_PhoPQ_compF1 and Prom_PhoPQ_FLAG_R1 primers (table S1).

### Sequence analysis and modelling

VgrG4 sequence (W9BAA4) was obtained from the UniProt Knowledgebase. A search for homologous crystal structures, which could be used as template structures, was performed with the Basic Local Alignment Search Tool (BLAST) against the Protein Data Bank using the VgrG4 sequence as query. Three structures were found: *P. aeruginosa* VgrG1 (PDB ID: 4uhv [41], 4mtk (Sycheva et al. to be published) and *E. coli* v3393 protein (PDB ID: 2p5z [84]). BLAST searches were also made in order to find sequences similar to both VgrG4 and PaVgrG1. Symmetry mates for PaVgrG1 (PDB ID: 4uhv [41]) were created in PyMOL in order to obtain the VgrG4 trimer. BLAST searches against the pdb database were likewise made using Sel1E (W9BPL7) sequence as queries. Only two structures (*P. aeruginosa* Pa5087 PDB ID: 5jkp and *P. aeruginosa* PA5087 PDB ID: 5jjo [85]) had an e-value < 0.001.

To determine the different domains of the protein, the sequences were analyzed with InterPro [43] and Prosite [47] and the secondary structures were predicted with PSIPred [86]. SignalP [46] was used to determine if the proteins contain any signal sequences and localization predictions were made with Psortdb [48], CELLO [49] and Phobius [50]. ConSurf [52] was used to analyze the evolutionary conservation. Pymol (The PyMOL Molecular Graphics System, Version 2.0 Schrödinger, LLC.) was used for visual inspection of the models and for creating pictures and the ABPS plugin was used to calculate the electrostatic potential of the protein surfaces.

The modelling of VgrG4 was performed in four separate parts. First, a structure-based sequence alignment was made for the N-terminal part (residues 1-563) using the crystal structures of PaVgrG1 (PDB ID: 4uhv [41], 4mtk (Sycheva et al. to be published)) and the *E. coli* v3393 protein (PDB ID:2p5z [84]). A few sequences similar in length to PaVgrG1 and to VgrG4, respectively, were aligned to the prealigned structural alignment to create the multiple sequence alignment for residues 1-563 (N-term), which was modelled using PaVgrG1 residues 1-549 as the structural template. Secondly, a region of the spike part (residues 564-636) in VgrG4 is predicted by PSIPred to contain two long alpha helices, which have no corresponding part in the PaVgrG1 protein. This part (Helix) was therefore modelled separately by the I-TASSER threading-method server [42], which uses multiple structural templates and builds a model by assembling fragments from the different templates. Thirdly, we made use of the repetitive pattern found in PaVgrG1 to model residues 637-677 (Repeat) by using a repeated part of the PaVgrG1 protein (residues 551-589) as structural template. Finally, residues 678-703 (C-term) were modelled based on the C-terminal part of PaVgrG1 (residues 550-645). 3D models for the N-terminal, Repeat and C-terminal parts were made with the homology modelling program Modeller [87], which created ten models, and the one with the lowest energy according to the Modeller objective function was then used for further analyses.

Sel1E was modelled with the threading server I-TASSER [42] and based on the homologs found in the BLAST search we added a restraint, which specified that I-TASSER would use the *P. aeruginosa* Pa5087 protein (PDB ID: 5jkp) as the template. The remaining part of the Sel1E protein, with no corresponding part in the template, were modelled independently by I-TASSER.

The models were visually inspected and evaluated by superimposition with the template structures using the program VERTAA in the Bodil modeling environment [88]. All models were also evaluated by the online model quality evaluation servers Qmean [89], PROSAweb [90], ProQ [91] and MODFOLD [51].

### Real time quantitative PCR to assess the expression of genes induced by ROS

Bacteria were collected in the exponential phase of growth, washed with PBS, and adjusted to an OD_600_ of 1 in PBS. 1 ml of this suspension was pelleted and resuspended in 1 ml of TRIzol (Invitrogen). RNA was purified following the indications of the manufacturer. The quality of the purified RNA was assessed using a nanodrop (NanoVue Plus). cDNA was generated from 1 μg of RNA using Moloney murine leukaemia virus (M-MLV) reverse transcriptase (Sigma-Aldrich) according to the manufacturer’s instructions. Quantitative real-time PCR analysis of gene expression was undertaken using the KAPA SYBR FAST qPCR Kit and Rotor-Gene Q instrument system (Qiagen). Thermal cycling conditions were as follows: 95 °C for 3 min for enzyme activation, 40 cycles of denaturation at 95 °C for 10 s and annealing at 60 °C for 20 s. Primers used in qPCR reactions are listed in Table S1. cDNA samples were tested in duplicate and relative mRNA quantity was determined by the comparative threshold cycle (ΔΔCT) method, using *rrsA* gene for normalisation.

### Fluorescence dye-based detection of ROS

*E. coli* MG1655 was grown until mid-exponential phase, bacteria were pelleted from 1-ml of the culture, washed once with PBS, and resuspended in 1-ml PBS. CM-H2DCFDA (Life Technologies, from 1 mM strock solution in 100% DMSO (vol/vol)) was added to bacteria to a final concentration of 10 μM. The mixture was incubated from 30 min at room temperature (20-22°C). One hundred microliters of the resultant cell suspension samples were transferred to a 96-well black plate, and fluorescence measured using a SpectraMax M2 Plate Reader (Molecular Devices) with excitation/emission wavelengths of 495/520 nm, and a slit width 1cm. Results are expressed as relative fluorescence units (RFU). All experiments were carried out with quintuple samples on at least three independent occasions.

### Single cell assessment of ROS by microscopy

*E. coli* MG1655 was grown until mid-exponential phase, and CM-H2DCFDA was added to the culture to a final concentration of 10 μM. One millilitre of the culture was spun down, washed twice with PBS and incubated in a 37 °C water bath for 30 min. 1 ml bacterial culture was then concentrated 10-fold by centrifugation, and 5 µl were spotted on a 1% agarose (w/v) pad containing 10 μM CM-H2DCFDA. After 30 min incubation at room temperature, fluorescence images were captured using the ×100 objective lens on a DM5500 microscope (Leica) equipped with the appropriate filter sets. Acquired images were analysed using the LAS imaging software (Leica).

For imaging of ROS in *E.coli* in contact with *K. pneumoniae*, Kp52.45 and *E. coli* MG1655 harbouring pJT04-*mCherry* were grown until mid-exponential phase, and CM-H2DCFDA was added to the *E. coli* culture to a final concentration of 10 μM. Bacteria were collected by centrifugation, washed with PBS, and mixed at 10:1 ratio (*Klebsiella:E. coli*). 1 ml of the sample were concentrated 10-fold by centrifugation, and 5 µl were spotted on a 1% agarose pad containing 10 μM CM-H2DCFDA. After 6 h incubation at 37°C, *Klebsiella* was labelled with rabbit anti-*Klebsiella* serum (1:1000) and visualised with goat anti rabbit Alexa Fluor 350 life-tech (1:1000) (Molecular Probes, A11046). Fluorescence images were captured using the ×100 objective lens on a Leica DM5500 microscope equipped with the appropriate filter sets. Acquired images were analysed using the LAS imaging software (Leica).

### Infection of Galleria mellonella larvae

*G. mellonella* larvae were obtained from UK Waxworms Limited. Upon receipt, larvae were stored in reduced light at 13°C with nil dietary supplementation. All experiments were performed within 7 days of receipt comprising larvae of 250–350 mg weight showing a healthy external appearance as previously described [39]. *K. pneumoniae* strains were prepared by harvesting refreshed 5 ml of exponential phase LB cultures (37°C, 180 rpm, 2.5 h), washing once in sterile PBS and then adjusting to an OD_600_ of 1.0 (i.e. ∼5 × 10^8^ CFUs/ml). Each suspension was thereafter diluted to the desired working concentration. Larvae were surface-disinfected with 70% (v/v) ethanol and then injected with 10 µl of working bacterial suspension at the right last proleg using a Hamilton syringe equipped with a 27-gauge needle. For each experiment, 10 larvae injected with sterile PBS were included as combined trauma and vehicle controls. Injected larvae were placed inside Petri dishes at 37°C in the dark. Larvae were considered dead when unresponsive to physical stimuli. Larvae were examined for pigmentation and time of death was recorded. The virulence assay was performed in triplicate (30 larvae total per strain).

### Cloning of Hcp1

*hcp1* was amplified by PCR using primers Hcp2467pBAD30_EcoRI_F1 and Hcp2468pBAD30_HindIII_R1 (Table S1), primer Hcp2468pBAD30_HindIII_R1 contains a VSV-G tag. The amplicon was gel-purified, digested with EcoRI and HinIII, gel-purified, and cloned into an EcoRI-HindIII-digested, Antarctic Phosphatase (New England Biolabs) treated pBAD30 expression vector to give pBADHcp1VSV. This plasmid was then transformed into *E. coli* β2163, and mobilized into *K. pneumoniae* strains by conjugation.

### Isolation of secreted Hcp1

Bacteria containing pBADHcp1VSV were grown until mid-exponential phase in 15-ml plastic tubes containing 5 ml of medium. Five millilitres of culture were centrifuged at 20,000 × g for 20 min and the supernatant was then filtered through a 0.2-μm filter. 450 μL of 100% trichloroacetic acid (TCA) solution were added to 4500 μL of the supernatant, and the sample was incubated overnight at −20°C. Proteins were recovered by centrifugation (15,000 × g for 10 min at 4 °C), and washed twice with ice cold acetone. The final precipitate was resuspended in 20 μL of SDS sample buffer.

### Immunoblotting

Proteins resolved by 15% SDS-PAGE and electroblotted onto nitrocellulose membranes. Membranes were blocked with 4% bovine serum albumin (w/v) in PBS and protein bands were detected with specific antibodies using chemiluminescence reagents and a G:BOX Chemi XRQ chemiluminescence imager (Syngene). The following antibodies were used: anti-Flag antibody (1:2,000; Sigma), anti-VSVG antibody (1:2,000; Sigma). Immunoreactive bands were visualized by incubation with horseradish peroxidase-conjugated goat anti-rabbit immunoglobulins (1:5000, BioRad 170-6515) or goat anti-mouse immunoglobulins (1:5000, BioRad 170-6516). To ensure that equal amounts of proteins were loaded, membranes were reprobed after stripping of previously used antibodies using a pH 2.2 glycine-HCl/SDS buffer with anti-*E.coli* RNA Polymerase α (1:2,000; BioLegend 663102).

GST fusions in yeast were immunodetected with primary rabbit polyclonal anti-GST (Z-5) antibody (1:2000, Santa Cruz Biotechnology). Secondary antibodies were IRDye-680 or IRDye −800 anti-rabbit and detection was performed with an Odyssey Infrared Imaging System (LI-COR Biosciences).

### Expression of vgrGs in S. cerevisiae

Yeast transformation was achieved by the standard lithium acetate protocol. Kp52145 *vgrGs* were cloned into the 2μ-based pEG(KG) vector to express GST fusion proteins in yeast [92] under the control of the GAL1 promoter. *vgrG1*, *vgrG2* and *vgrG4* were amplified by PCR using primers carrying BamHI and XbaI restriction sites for *vgrG1*, and XbaI restriction sites for *vgrG2* and *vgrG4* (Table S1). All PCR products were cleaved with the corresponding enzymes and inserted into the same site in the pEG(KG) vector and sequenced. *vgrG4* truncated variants were obtained by sit-directed mutagenesis using primers indicated in Table S1.

*S. cerevisiae* strains expressing VgrGs were grown in Synthetic complete medium (SC) containing 0.17 % (w/v) yeast nitrogen base without amino acids, 0.5 % (w/v) ammonium sulphate and 2 % (w/v) glucose, and was supplemented with appropriate amino acids and nucleic acid bases. SCGal and SCRaf media were SC with 2 % (w/v) galactose or raffinose, respectively, instead of glucose. Galactose induction experiments in liquid media were performed by growing cells in SCRaf medium at 30 °C to exponential phase and then adding galactose to 2 % (w/v) for 4-6 h. Effects of the expression of VgrG genes on yeast growth were tested by spotting cells onto SC and SCGal plates lacking the corresponding auxotrophic marker to maintain the plasmid. Briefly, transformants bearing pEG(KG)-based plasmids were grown overnight in SC lacking uracil and leucine (SC–Ura-Leu) and adjusted to an OD_600_ of 0.5. This and three serial 1/10 dilutions were spotted on the surfaces of SC or SCGal solid media lacking uracil. Growth was monitored after 2–3 days at 30°C.

### Statistical analyses

Statistical analyses were performed using two-way ANOVA with Bonferroni correction for multiple comparisons, or unpaired T-test. Survival analyses were undertaken using the Log-rank (Mantel-Cox) test with Bonferroni correction for multiple comparisons (α=0.008). All analyses were performed using GraphPad Prism for Windows (version 5.03) software. P-values of < 0.05 were considered statistically significant.

## ACKNOWLEDGEMENTS

We thank the members of the J.A.B. laboratory for their thoughtful discussions and support with this project. We also thank the bioinformatics (J.V. Lehtonen), translational activities and structural biology infrastructure support from Biocenter Finland and Instruct-FI, and CSC IT Center for Science for computational infrastructure support at the Structural Bioinformatics Laboratory, Åbo Akademi University. D.S. is the recipient of a PhD fellowship funded by by the National Doctoral Programme in Informational and Structural Biology, Svenska Kulturfonden and Orion Research Foundation funding to M.Å., by Sigrid Jusélius Foundation and Tor, Joe, and Pentti Borg’s Foundation funding to T.A.S. This work was supported by Biotechnology and Biological Sciences Research Council (BBSRC, BB/N00700X/1, and BB/L007223/1) and Queen’s University Belfast start-up funds to J.A.B. The funders had no role in study design, data collection and analysis, decision to publish, or preparation of the manuscript.

